# The Chromosome-level Genome of *Dracaena cochinchinesis* Provides Insights into its Biological Features and the Mechanism of Dragon’s Blood Formation

**DOI:** 10.1101/2022.02.08.479636

**Authors:** Yanhong Xu, Kaijian Zhang, Zhonglian Zhang, Yang Liu, Feifei Lv, Peiwen Sun, Shixi Gao, Qiuling Wang, Cuicui Yu, Jiemei Jiang, Chuangjun Li, Meifang Song, Zhihui Gao, Chun Sui, Haitao Li, Yue Jin, Xinwei Guo, Jianhe Wei

## Abstract

*Dracaena*, a remarkably long-lived and slowly maturing species, is famous all over the world for the production of dragon’s blood, a precious traditional medicine used by different cultures since ancient times. However, lacking a high-quality genome,the molecular mechanisms underlying these traits are largely unknown and that greatly restricts the protection and regeneration of the rare and endangered plant resources. Here, we sequenced and assembled a chromosome-level genome of the *Dracaena cochinchinensis*, the first to be sequenced of *Dracaena* Vand. ex L. The *D. cochinchinensis* genome covering 1.21 Gb with a scaffold N50 of 50.06 Mb, and encodes 31,619 predicted protein-coding genes. We found *D. cochinchinensis* has undergone two whole genome duplications (WGDs) and two long terminal repeats (LTRs) insertion burst events. The expansion of *cis*-zeatin O-glucosyltransferase (*cZOGT*) and small auxin up-regulated RNA (*SAUR*) genes is account for its longevity and slow growth. In flavonoids biosynthesis pathway, transcription factors bHLH and MYB were predicted as the core regulators, and ROS as the specific signal molecule during the process of injury-induced dragon’s blood formation. Our study not only provides high-quality genomic knowledge of *D. cochinchinensis*, but also deciphered the mystery of its longevity, and preliminarily elucidated the molecular mechanism of dragon’s blood formation, which will facilitate the resource protection and sustainable utilization of *Dracaena*.

**SHORT SUMMARY:** This study reports a chromosome-level genome assembly for *D. cochinchinensis*, the first genome of *Dracaena* Vand. ex L. It 29 provides valuable genetic resources and creates a large scope for studying *Dracaena*. We found the significant expansion of genes associated with its longevity and slow growth, and genes in flavonoids biosynthesis were first completely identified. Moreover, transcription factors bHLH and MYB as the core regulator of flavonoids biosynthesis, and ROS as the specific signal molecule during the process of injury-induced dragon’s blood formation were also identified. These results not only deciphered the mystery of its longevity, but also elucidated the molecular mechanism of dragon[s blood formation preliminarily, which will facilitate the resource protection and sustainable utilization of *Dracaena*.

## INTRODUCTION

Dragon’s blood, the red resin produced by tree species in the genus *Dracaena* (Asparagaceae), is a precious traditional Chinese medicine, and a folk medicine used by many nationalities in the world. It has several therapeutic functions of activating blood circulation and dissipating blood stasis, relieving inflammation and pain, astringency and hemostasis, etc (Gupta *et al*., 2008; Fan *et al*., 2014; Sun *et al*., 2019). Besides being a famous medicine, it is also used as the colorant of art works by many cultures (Zheng *et al*., 2012). It is called “dragon’s blood” because the resin or sap excreted by a part of the genus *Dracaena* plants is deep red and looks like the blood of a legendary dragon when encountered external injury or microbial invasion (Gupta *et al*., 2008), and therefore, *Dracaena* has been famous all over the world for its red resin, and belongs to the cultural heritage of humanity.

*Dracaena* species are monocotyledonous evergreen plants mainly distributed in tropical and subtropical regions of Asia and Africa. They are also a remarkably long-lived and slowly-growing species that known as “plant longevity star” (Zheng *et al*., 2012; Tomlinson & Huggett, 2012; Maděra *et al*., 2018, 2020; Scheiger *et al*., 2020). The age of a fabulous dragon blood tree was overestimated for several thousands of years (Humboldt, 1814; Christ *et al*., 1886; Schenck, 1907). Thereafter, through the observation of a few largest trees, it was concluded that their ages are about several hundred years, and the flowering time varies from ten to several decades (Pütter, 1925; Symon, 1974; Mägdefrau, 1975; Zheng *et al*., 2012; Maděra *et al*., 2018). According to the research of Maděra (2018), the juvenile stages of *D. cinnabari* from seedling to the first flowering will take about 200-250 years, even more (Maděra *et al*., 2018), demonstrating it is an extremely slowly growth and slowly maturing species. Due to the difficulties in the authentic study of natural population aging, several methodologies of age estimation were developed and improved (Adolt & Pavliš, 2004; Adolt *et al*., 2012; Jura Morawiec, 2019; Lengálová *et al*., 2020). Anyway, they all showed that the dragon tree is a long-lived species.

The origin and identity of dragon’s blood have been confused, and there are many source plants of dragon’s blood, such as *Croton, Dracaena, Daemonorops* and *Pterocarpus* (Gupta *et al*., 2008; Ding *et al*., 2020). In nature, *Dracaena* grows slowly, and only trees over 30-years-old have the possibility to produce dragon’s blood (Edwards *et al*., 1997; Wang *et al*., 2013; Ding *et al*., 2020). For a long time, due to the continuous over-exploitation, its natural resources are threatened with depletion. In 1998, many of the *Dracaena* species were list in the International Union for Conservation of Nature Red List (IUCN red list, 2017.2). Of them, *D. cochinchinensis* and *D. Cambodian* species in China were rated as vulnerable (Hubálková *et al*., 2015; Kamel *et al*., 2018). Soon afterwards, the two species had been listed in the List of National Key Protected Wild Plants of China, which are the second grade of national key endangered protection and prohibited to felling. For the endangered status coupled with limited genetic resources, it is critical to consider adequate measures for conservation management and the exploitation of dragon’s blood.

In China, *D. cochinchinensis* is the official original species produced dragon’s blood (Zheng *et al*., 2009; Luo *et al*., 2011), which was first discovered by Xitao Cai in 1972 in Yunnan province and has been used to be the substitute of the traditionally imported dragon’s blood, called Long-Xue-Jie (Chinese dragon’s blood). According to literature research, the dragon’s blood has been recorded and applied in China for at least 1500 years, and the authentic source of Chinese ancient *Dracaena* should be imported from Africa (Cai & Xu, 1979). With the rising of shipping, the genuine *Dracaena* has changed from the resin of *Dracaena* in agave family to the red resin of the fruit of *rattan* in Southeast Asia until the discovery of *D. cochinchinensis* which is exclusively distributed in China (Cai & Xu, 1979; Zheng *et al*., 2006; Wang *et al*., 2011).

Flavonoids are the main components of dragon’s blood. Flavonoid oligomers, composing of one dihydrochalcone unit condensed with one or more chalcone, flavane or homoisoflavane units, constituted the major identified components of dragon’s blood from the genus *Dracaena* (Pang et al., 2016). The oligomeric flavonoids account for over 50% of the resin from *Dracaena* spp. by weight (Hao et al., 2015b). Up to now, hundreds of flavonoids have been isolated from the resin wood of *D. cochinchinensis* (for review Gupta *et al*., 2008; Fan *et al*., 2014, Sun *et al*., 2019). Additionally, terpenoids and steroidal saponins are also effective components of dragon’s blood (Fan *et al*., 2014; Teng *et al*., 2015; Maděra *et al*., 2020; Vanickova *et al*., 2020), which were reported to be cytotoxic and could be used as an anticarcinogen (Darias *et al*., 1989; Mimaki *et al*., 1999; Gonźalez *et al*., 2003; Herńandez *et al*., 2004; Ding *et al*., 2020; Maděra *et al*., 2020).

Though there are many studies on the chemical constituents and pharmacological effects of dragon’s blood (for review Gupta *et al*., 2008; Fan *et al*., 2014; Sun *et al*., 2019; Maděra *et al*., 2020), few about its formation mechanism and the molecular basis about its biology, even many knowledge gaps in the research of *Dracaena*. In this study, we first report the genome draft of *D. cochinchinensis* at chromosome level, as well as the transcriptome and metabolome analysis were conducted, which is undoubtedly an important step to understand the genetic and molecular basis of its fundamental biology and conservation management.

## RESULT

### Genome Sequencing, Assembly and Annotation

The genome survey using a k-mer analysis (k = 17) revealed that the genome size of *D. cochinchinesis* was approximately 1.257 Gb, with the heterozygosity of 0.52% and the repeat sequence ratio of 67.8% (Fig. S1; Table S1). Based on the estimated genome size, using a combination of SMRT sequencing technology from PacBio, 10xGenomics and short-read sequencing from Illumina platforms, we totally obtained 407.06 Gb clean data, which represented x 323.75 coverage (Table S2). The initial assembly is 1.21 Gb, containing 1898 scaffolds and 3042 contigs, with a scaffold N50 size of 2.07 Mb and contig N50 size of 1.1 Mb (Table S3). Assessment of the completeness of the genome assembly with CEGMA indicated 96.77% coverage of the conserved core eukaryotic genes, and Benchmarking Universal Single-Copy (BUSCO) (Simao *et al*., 2015) analyses showed that the genome was 93.6% complete (**Table S4**), suggesting that the genome assembly was relatively complete and of high quality. Next, ∼137.26 Gb high-quality Hi-C data were used to further improved the genome assembly, and 90.26% of the genome were anchored onto 20 chromosomes (2n=40) (**Fig. 1b; Fig. S2**;**Table S5**). The final assembly consists of 1835 scaffolds, which spans 1.21 Gb, with a scaffold N50 size of 50.05 Mb (**Table S6**). The *D. cochinchinesis* chromosomes showed a significant level of synteny with its related species *A. officinalis* (**Fig. S3)**.

**Fig. 1.**
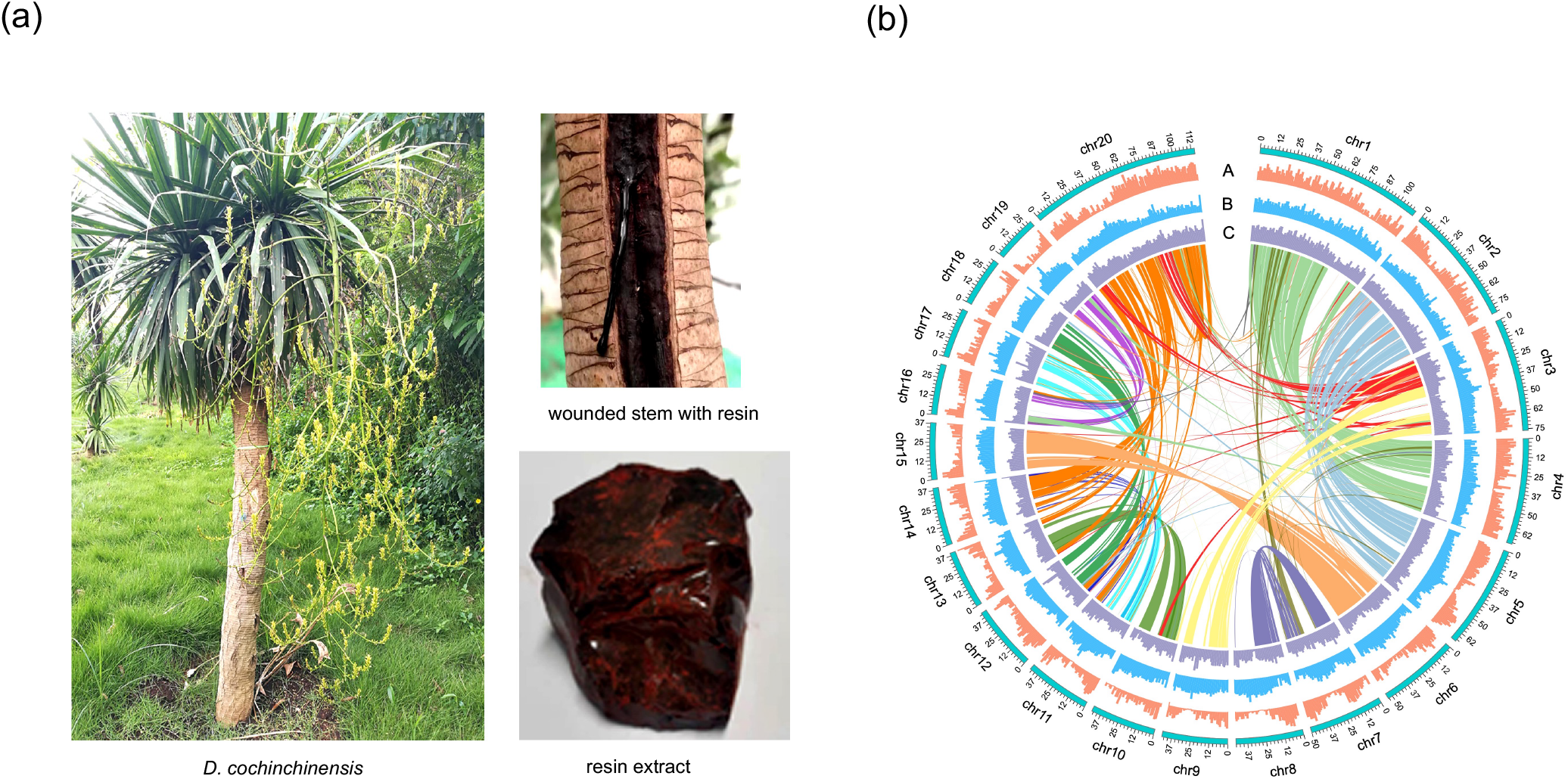
Morphology and genome features of *D. cochinchinensis*. (a) morphology of *D. cochinchinensis* and dragon’s blood. (b) Genome features and synteny information. The circus map, from outside to inside, shows GC density, repeat density and gene density, respectively.

Based on *de novo* and homology-based predictions and transcriptome data, a total of 31,619 protein-coding genes were predicted, and 31,402 genes (99.3%) were assigned with functions (**Table S7**). In addition, by comparing with the known ncRNA libraries, the ncRNA information of the *D. cochinchinesis* genome of was obtained (**Table S8**), including 1570 transfer RNAs (tRNAs), 1278 ribosomal RNAs (rRNAs), 1576 microRNAs (miRNAs) and 722 small nuclear RNAs (snRNAs).

### Genomic Evolution of *D. cochinchinesis*

*D. cochinchinesis* belongs to the Asparagales clade of monocots. It is close to *A. officinalis, D. officinalis* and *A. shenzhenica* that have completed genome sequencing. We clustered the annotated genes into gene families among them plus other additional monocotyledon and dicotyledon species, includ*ing E. guineensis, O. sativa, A. comosus, Arabidopsis, Populus, A. sinensis, H. brasiliensis, C. grandis, C. papaya, E. grandis, V. vinifera, A. Trichopoda*. Family clustering yielded 31,324 gene families (groups), including 252 single-copy families. CDS-based evolutionary trees were constructed using RAXML software. *D. cochinchinensis* and *A. officinalis, D. officinalis* and *A. shenzhenica* are gathered in one branch, respectively, which indicated that *Asparaginaceae* and *Orchidaceae* had an independent diverge long ago (138.4 Mya), while diverge time between *D. cochinchinensis* and *A. officinalis* is 87.7 Mya (**Fig. 2a**).

**Fig. 2.**
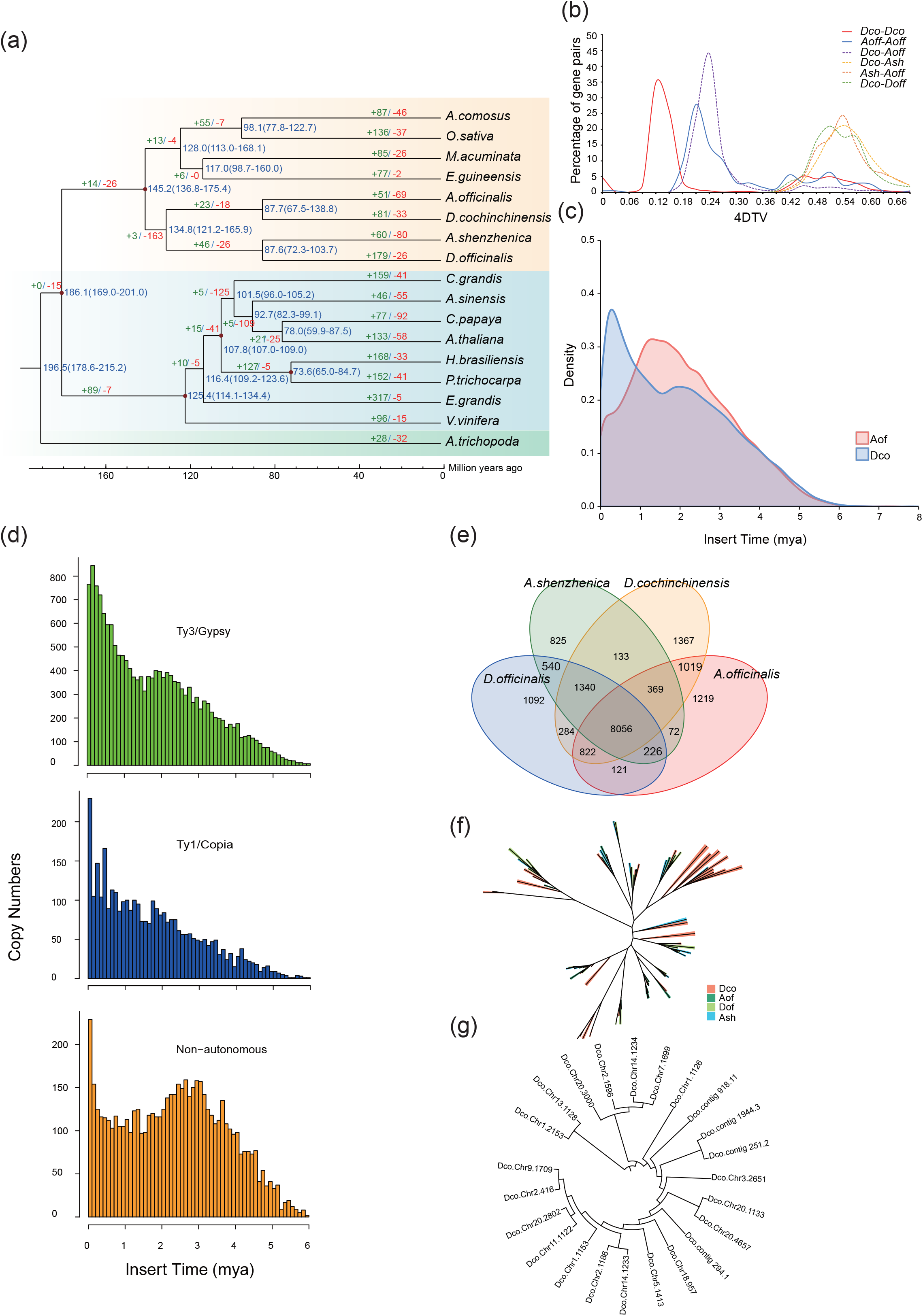
Comparative Genomic Analysis. (a) Phylogenetic tree of 17 plant species. Divergence time (Mya) estimations are indicated by the blue numbers beside the branch nodes. The number of gene-family contraction and expansion is indicated by green and red numbers respectively. (b) Distribution of 4DTv for paralogous genes among *D. cochinchinensis, A. officinalis, D. officinalis* and *A. shenzhenica*. (c) Estimation of LTR activity showed there are two insertion burst events in *D. cochinchinesis* genome. (mya). (d) In this recent burst of activity, the majority of elements responsible were both autonomous (3359 copies of Ty1-copia and 16897 copies of Ty3-gypsy) and non-autonomous LTRs (5861 copies), with peaks of activity <6 mya. (e) Venn diagram shows shared and unique orthologues among the four asparagus species, including *D. cochinchinesis, A. officinalis, D. officinalis* and *A. shenzhenica*, and genes unique to *D. cochinchinesis*. (f-g) Phylogeny of the SAURs and ZOGTs family in *D. cochinchinensis*.

To investigate the genome expansion in *D. cochinchinensis*, we analyzed whole-genome duplication (WGD) events. 4DTv values was estimated on the basis of the paralogous gene pairs in collinear regions detected in *D. cochinchinensis* and three other representative plant species, *A. officinalis, D. officinalis* and *A. shenzhenica*. The distribution of 4DTv distances in *D. cochinchinensis* showed two peaks at the values of approximately 0.12 and 0.45 (**Fig. 2b**). The first peak shared by *D. cochinchinensis* and *A. officinalis* may correspond to the Asparagales-β event previously identified in the *A. officinalis* genome (Harkess *et al*., 2017; Li *et al*., 2020). The second peak at approximately 0.12 indicate that *D. cochinchinensis* underwent another WGD event in dependently after diverging from *A. officinalis* (**Fig. 2b, Fig. S4**). The 29,280 duplicated genes were classified into five different categories: 6864 whole-genome duplicates (WGD duplicates, 23.1%), 1506 tandem duplicates (TD, 5.1%), 1196 proximal duplicates (PD, 4.1%), 7301 transposed duplicates (TRD, 24.9%), and 19613 dispersed duplicates (DSD, 66.7%). We further investigate the duplications of genes involved in flavonoids biosynthesis (i.e., *PAL, 4CL, C4H, CHI, CHS, DFR, F3H, FLS, FAR, PPO, OMT*), and found some genes were differently duplication, and there were more genes retained by WGD, suggesting the recent WGD event was important to the evolution of flavonoid biosynthesis in *D. cochinchinensis* (**Table S9**).

60.15% (729.11 Mb) of the *D. cochinchinesis* genome comprises repetitive sequences (**Table S10**), similar to *D. officinale* (63.33%) (Yan *et al*., 2015) and *E. guineensis* (57%) (Singh *et al*., 2013), but higher than *A. shenzhenica* (42.05%) (Zhang *et al*., 2017) and *A. comosus* (43.7%) (D’Hont *et al*., 2012). As in many other sequenced plant genomes (Kim *et al*., 2014; Teh *et al*., 2017; Zhang *et al*., 2017), long terminal repeats retrotransposons (LTR-RTs) were the dominate, accounts for 51.06% of the assembly, with the *Gypsy* (501,175,881bp, 41% of the genome sequence) and *Copia* (72,176,608 bp, 5.9% of the genome sequence). Furthermore, analyzing the insertion time of intact LTR, it is found there are two insertion burst events in *D. cochinchinesis* genome occurred recently and before, about 0.3-0.4 million years and 2 million years, respectively (Fig. 2c), dominated by both autonomous and non-autonomous LTRs (Fig. 2d). In order to investigate the evolution of transposable elements (TEs) in *D. cochinchinesis*, phylogenetic trees of domains in reverse transcriptase genes were constructed for both *Copia* and *Gypsy* superfamily. We found the majority of *Gypsy* was species-specific in *D. cochinchinesi*s, while the *Copia* superfamily displayed a different pattern with consisting of elements from all three species (**Fig. S5**).

### Comparative Genomic Shows the Adaption of *D. cochinchinesi*s

Copy numbers of homologous gene families vary greatly among different species, which is caused by the differences in the rates of gene gain and loss. Among the 31,318 gene families inferred to be present in the most recent common ancestor of the 17 species that used for gene family analysis, 81 gene families were found to have undergone expansion in *D. cochinchinensis* relative to the other 16 species (**Fig. 2a**). A total of 730 and 1088 genes in the expanded families were annotated to Gene Ontology (GO) terms and Kyoto Encyclopedia of Genes Genomes (KEGG) pathways, respectively. GO analysis revealed that these expanded orthogroups were related to several metabolism process (**Table S11, Figs, S6-7**), such as “ATP metabolic process”, “defense response” based on the biological process category, and “binding”, “catalytic activity”, “oxidoreductase activity”, “peroxidase activity” based on molecular function category. The enriched KEGG categories were the categories of “plant-pathogen interaction”, “phenylpropanoid biosynthesis”, “zeatin biosynthesis”, “sesquiterpene and triterpenoid biosynthesis”, “flavone and flavonol biosynthesis”, and “photosynthesis”, etc (**Table S12, Fig. S8**). We speculate that these expanded genes are closely related in some way to the function requirements of environmental adaptability of *D. cochinchinensis* and the formation of dragon’s blood.

It is noted that 28 genes were found to be under positive selection, which significantly enriched in KEGG terms related to base excision repair (BER), nucleotide excision repair (NER), mismatch repair (MMR) and DNA replication (**Table S13**). They include DNA ligase 1 and DNA excision repair protein which are DNA-repair pathways that play crucial roles in maintaining genomic integrity particularly in response to exposure to environmental genotoxicants including ultraviolet and reactive oxygen produced from the harsh rays of the sun (Schuch *et al*., 2017; Nisa *et al*., 2019). The response of cells to DNA damage will restore the DNA structure to its original function, or just enable the cell to survive the damage.

We further identified unique and shared gene families among the four Asparagus species, and found 8056 were shared as families with *A. officinalis, D. officinalis* and *A. shenzhenica*, and 1367 gene families, containing 2954 genes appeared to be unique (**Fig. 2c; Tables S14-15**). Functional annotation revealed that two classes of genes, small auxin up-regulated RNA (*SAUR*) genes and *cis*-zeatin O-glucosyltransferase (*cZOGT*) in zeatin biosynthesis, may be closely related to its slow growth and longevity. There are totally 40 *SAURs* identified in *D. cochinchinensis*, while 20 in *D. officinalis*, 14 in *A. officinalis*, 18 in *A. shenzhenica*, and 23*cZOGTs* is unique to *D. cochinchinensis*, which was not identified in the three closely related species (**Fig. 2f-g**). SAUR is an auxin-responsive protein that acts as the plant’s toolbox for adaptation of growth and development (Ren & Gray, 2015; Stortenbeker & Beme, 2019). Studies in rice confirmed SAUR inhibit growth via negatively regulates auxin synthesis and transport, in addition, SAUR significantly increased anthocyanin content and the ABA level (Kant *et al*., 2009). *ZOGT* are known led to growth retardation and senescence delay (Rodo *et al*., 2008; Kudo *et al*., 2012). Therefore, the expansion of *SAURs* and *cZOGTs* is predicted to at least partly account for its slow growth and long life, even its adaptation to the arid environment of karst landform.

### Metabolic Profiling and Differential Metabolite Pathways in the Process of Dragon’s Blood Formation

Dragon’s blood does not appear directly after the wound, the process of its formation takes time. From the slices, we observed the color of resin gradually deepened during the formation of dragon’s blood, with filling the vascular bundle and surrounding parenchymal cells after 90 d (**Fig. 3a**). Eventually, the resin might be in the form of tear-like drops or chips which coats the place of wound (**Fig.1a**), protecting the plant from the spread of pathogens. Thus, it is the product of dragon tree defense mechanism (Wang, 2007; Jura-Morawiec & Tulik, 2016; Maděra *et al*., 2020).

**Fig. 3.**
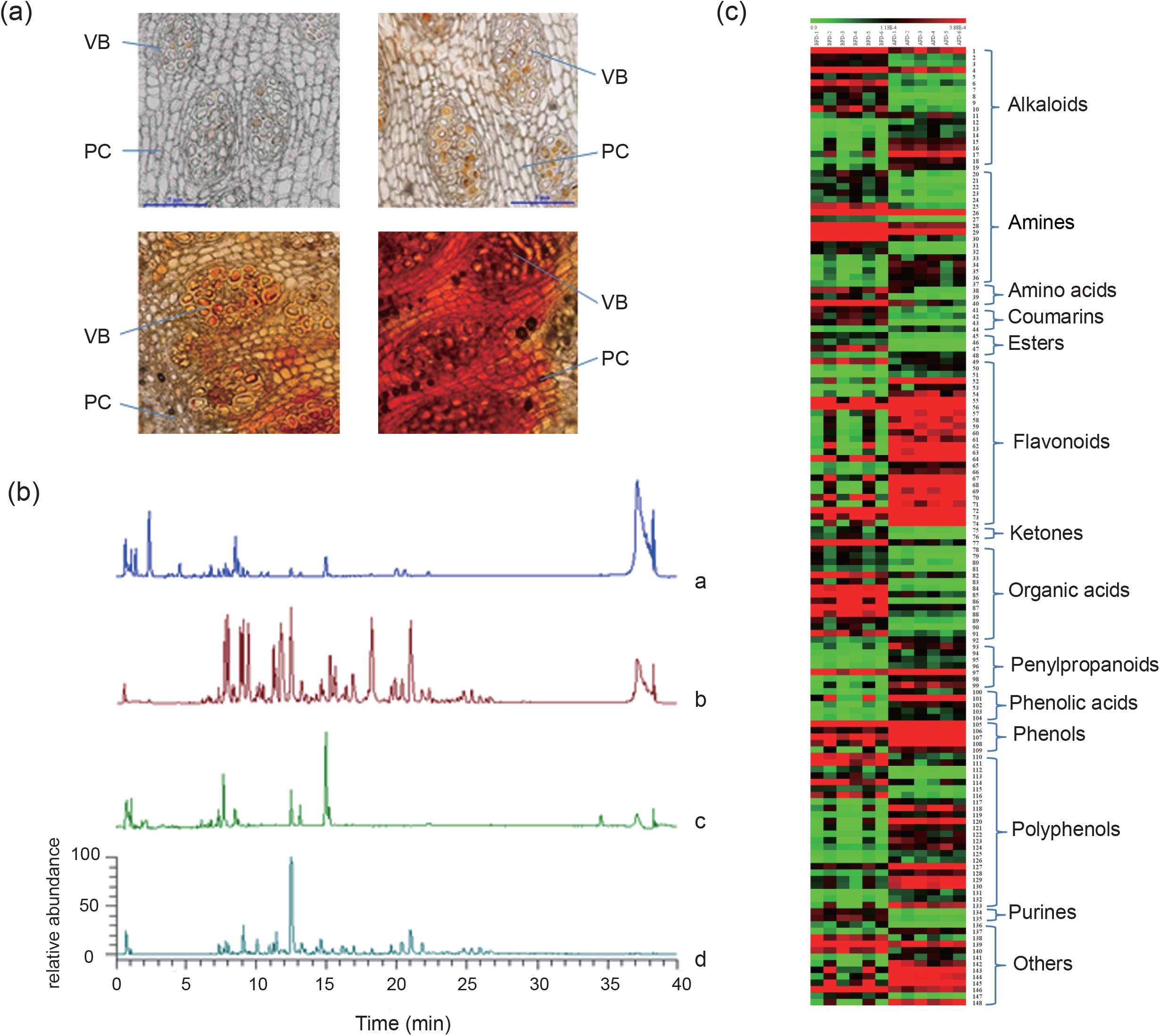
Histological and Metabolic Profilings during dragon’s blood formation. (a) Histological images during the formation of dragon’s blood in *D. cochinchinensis*. (The yellow and red substances in the picture are dragon’s blood resin. VB = Vascular Bundle, PC: parenchymal cell.) (b) Typical total ion chromatograms of *D. cochinchinensis*. (a: positive TIC before dragon’s blood formation; b: positive TIC after dragon’s blood formation; c: negative TIC before dragon’s blood formation; d: negative TIC after dragon’s blood formation.) (c) Heat map of identified metabolites of *D. cochinchinensis* before and after the formation of dragon’s blood (BFD 1∼6 represents six duplications of the analytical materials before formation of dragon’s blood; AFD1∼6 represents six duplications of the analytical materials after formation of dragon’s blood).

To elucidate the changes in metabolite composition during the formation of dragon’s blood, we focused on identifying metabolites with significant difference by filtering the picked peak. Ultimately, a total of 148 differential metabolites were identified or tentatively identified (**Table S16; Fig. 3c**). As shown in Fig. 3c, content of most compounds changed significantly after injury. Among them, 85 increased, including flavonoids, phenols, polyphenols, penylpropanoids and phenolic acids, and 63 compounds, mainly organic acids, amines and steroid were decreased. Consistently, the total normalized peak areas showed the same results (**Fig. S9**).

Biological processes associated with the formation of dragon’s blood were investigated using transcriptome data. The up-regulated differentially expressed genes (DEGs) are mainly enriched in “metabolic pathway”, “biosynthesis of secondary metabolites”, showing that different time points correspond to different metabolic processes (**Fig. 4**). For example, the first metabolic pathways to change in 2 h are plant-pathogen interaction, plant hormone signal transduction as well as starch and sucrose metabolism. Then, biosynthesis of amino acid and citrate cycle were significantly enriched, and flavonoid biosynthesis appeared at 3 d and lasted until 17 M, which was the most active metabolic pathway, and terpenoid backbone biosynthesis at 15 d, sesquiterpenoid and triterpenoid biosynthesis at 30 d, phenylpropanoid biosynthesis at 6 M. The differentially expressed metabolic pathways combined with the metabolic profiling showed in Fig. 3c, we infer that the synthesis of dragon’s blood involves a series of very complex signal transduction, gene expression regulation and metabolism processes.

**Fig. 4.**
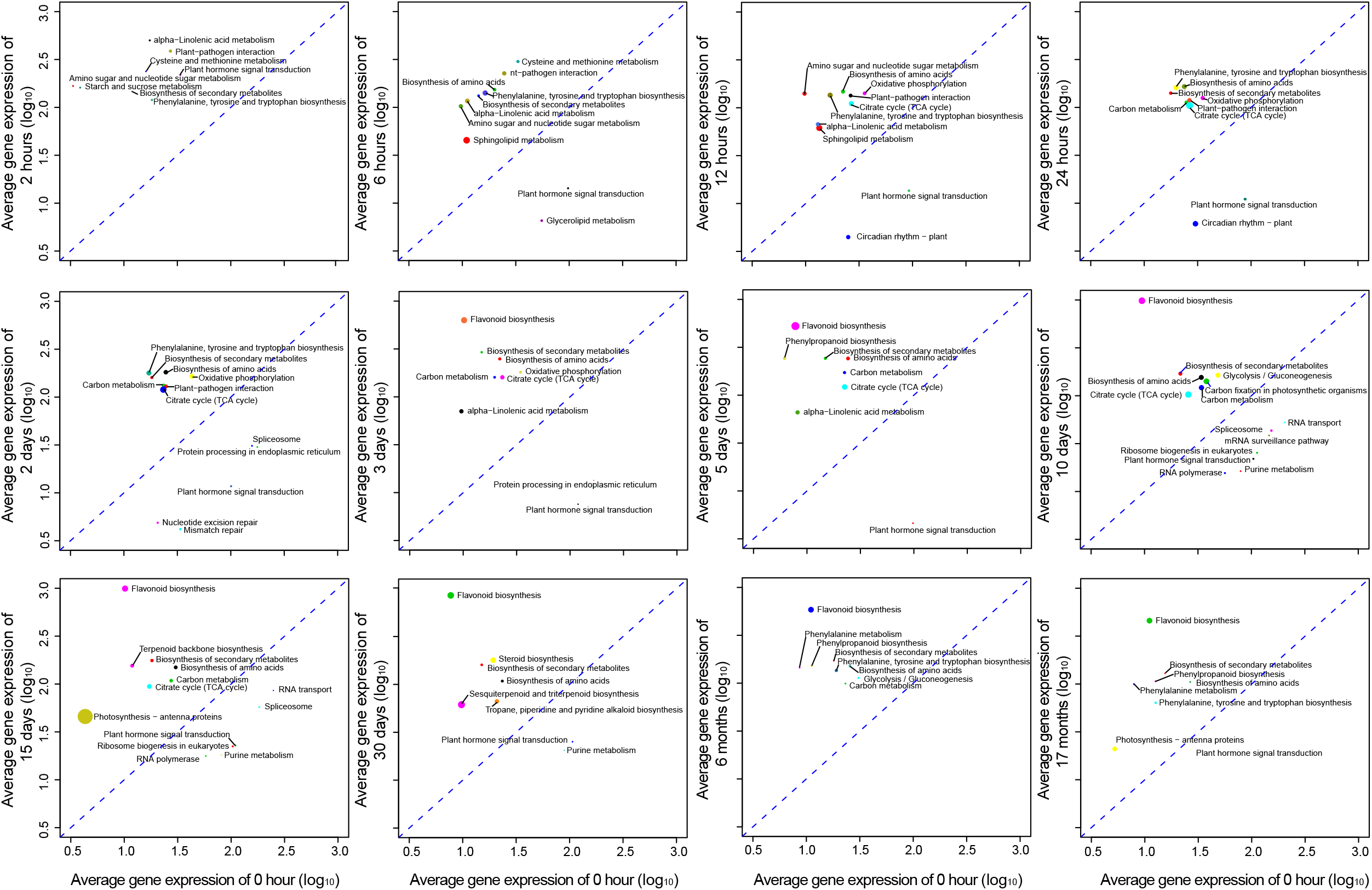
Differentially expressed metabolic pathways during dragon’s blood formation. Transcriptome analysis identifies differentially expressed pathways during dragon’s blood formation. The up-regulation pathway was above the oblique line, and the down-regulation pathway was below the oblique line. The circle size represents the number of genes, and the coordinate value represents the average up-regulation or down-regulation times.

### Genes in Flavonoids Biosynthesis in *D. cochinchinensis*

Numerous studies have shown that flavonoids are the main components of dragon’s blood (Gupta *et al*., 2008; Fan *et al*., 2014). It occurs via phenylpropane biosynthesis pathway (**Fig. 5**). From our genome assembly, 11 gene families related to dragon’s blood formation were identified (**Table S17**), including some common enzymes in the flavonoid synthesis pathway: PAL, C4H, 4CL, CHS, CHI, F3H, FLS, DFR, and downstream synthetic genes producing specific components of dragon’s blood: such as OMT (O-methyltransferase) to form loureirin A and PPO (polyphenol oxidase) to form loureirin B, which is the index components of the standard of dragon’s blood in China. Gene expression results showed that four of the five *CHS* genes and two *CHI* genes showed significantly upregulated after injury and lasted until 17 M. Similarly, two of the three *F3H* genes are not expressed in healthy tissues, and the highest expression level is at 3 d and 5 d after injury. Some of *FLS* and *PPO* members are also almost not expressed in health stem, but expressed after injury (**Fig. 5**). We speculated that these genes may be specifically involved in the formation of dragon’s blood, and their expression is significantly induced by injury.

**Fig. 5.**
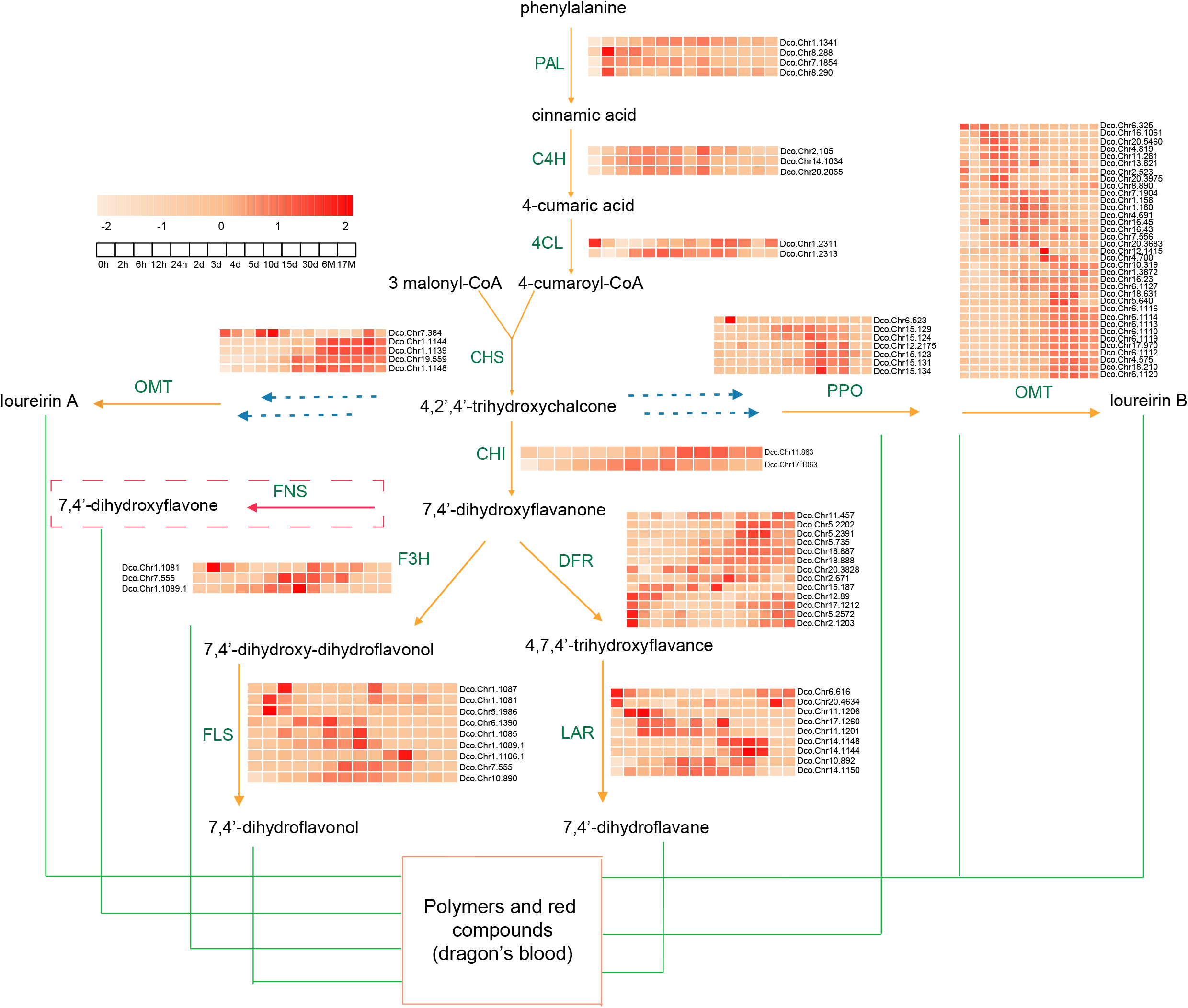
Flavones biosynthesis in *D. cochinchinensis*. The flavones biosynthesis pathway and expression profiles (reads per kilobase per million reads mapped; RPKM) of the genes involved in rubber biosynthesis. PAL, phenylalanine ammonia-lyase; C4H, cinnamate 4-hydroxylase; 4CL, 4-coumarate CoA ligase; CHS, chalcone synthase; CHI, chalcone isomerase; FNS, flavone synthase; F3H, flavanone 3-hydroxylase; DFR, dihydroflavonol 4-reductase; FLS, flavonol synthase; LAR, leucoanthocyanidin reductase; PPO, polyphenol oxidase; OMT: O-methyltransferase.

Further, based on co-expression model, two kinds of transcription factors bHLH and MYB, expressed in the wounded stems, were predicted as the core regulators in the synthesis of flavonoids, which are likely target to *PAL, CHS* and *FLS* and participate in the regulation of flavonoid synthesis pathway (**Fig. S10; Table S18**). Analyzing the 2000 bp upstream of the gene coding region, we found many *cis*-acting elements specifically recognized by bHLH and MYB (**Table S19**), which further demonstrated these enzyme genes are likely regulated by the two groups of transcriptional factors. Moreover, from the expression heat map of bHLH and MYB (**Figs. S11-12**), we can see that they are significantly induced after injury 2 h. We speculate that bHLH and MYB not only regulates a key enzyme gene *PAL, CHS* and *FLS*, but also regulates the whole flavonoids synthetic pathway.

### Specific Regulation of ROS Signaling during Dragon’s Blood Formation

Focused on analysis genes involved in different signaling pathways showed that genes related to ROS production and scavenging is substantially up-regulated (**Fig. S13**). Oxidative burst is a common early event in plant defense responses. ROS will accumulate rapidly when plants are injured outside. ROS generation depends on NADPH oxidase and apoplastic peroxidases (Bolwell & Wojtaszek, 1997), and peroxisomes are probably considered the major sites of intracellular H_2_O_2_ accumulation as a consequence of their oxidative metabolism (del Río *et al*., 2006). From the transcriptome data, we found that several enzymes, such as respiratory burst oxidase homolog (RBOH), amine oxidases (AOC), NAD(P)H oxidases (NOX), peroxidase (PRX), which contribute to ROS generation were significantly upregulated at 2 h after injury (**Fig. S13**). Meanwhile, ROS scavenging enzymes, such as superoxide dismutase (SOD), ascorbate peroxidase (APX), glutathione peroxidase (GPX) and catalase (CAT), are also up-regulated subsequently (**Fig. S13**). Obviously, the degree of oxidative stress is determined by the level of ROS, and the balance between ROS and antioxidants is essential to maintain balanced redox state.

H_2_O_2_ is the most stable type of ROS within plants. To preliminarily validate the H_2_O_2_ response during the dragon’s blood, the change of H_2_O_2_ content within stem of *D. cochinchinensis* over time after mechanical damage is detected. It was showed H_2_O_2_ concentration obviously improved 1 h after mechanical damage, and continued to increase until the maximum value at 6 h. After 8 h, it began decreased slightly until the minimum value was reached at 5 d (**Fig. S14**). In contrast, other signals, such as ethylene, nitric oxide and brassinosteroids, have no obvious changing rule (**Figs. S15-17**). We infer ROS as the first sensor and the core regulator in the process of dragon’s blood formation (**Fig. 6**), which is consistent with the conclusion that ROS represent a point at which various signaling pathways come together (Camejo *et al*., 2016; Sewelam *et al*., 2016),

**Fig. 6.**
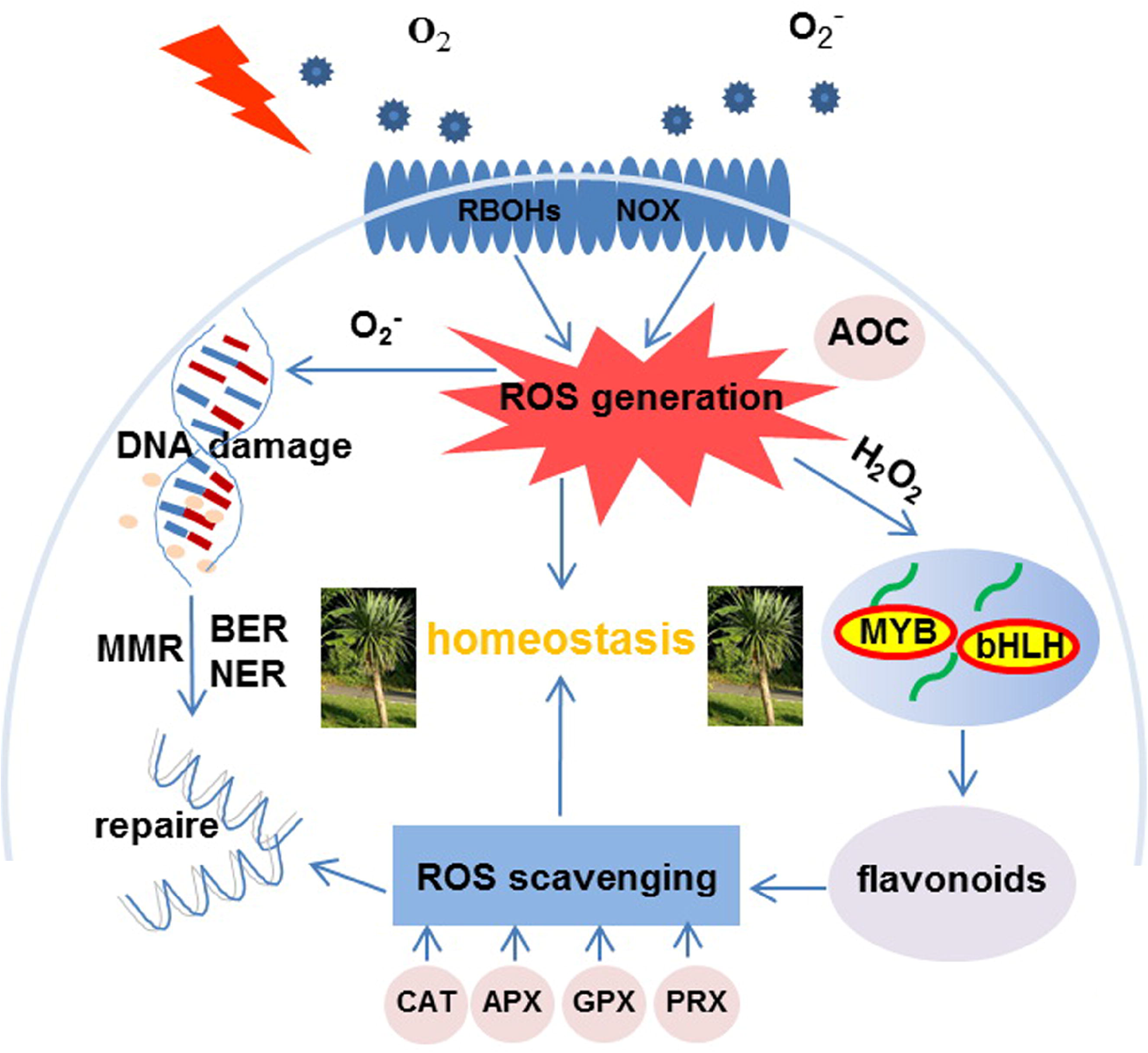
A propose model of ROS as a specific regulating signal in the adaptation of *D. cochinchinensis*. External wound leads to ROS burst and excessive ROS causes DNA damage. Fortunately, most of the positive selection genes in the genome are enriched in DNA-repair pathways, such as base excision repair (BER), nucleotide excision repair (NER), mismatch repair (MMR),which play crucial roles in maintaining genomic integrity. Meanwhile, ROS as a signal molecule activates transcription factors, thus activating the flavonoid synthesis pathway to synthesize a large amount of flavonoids. Flavonoid is the main component of dragon’s blood, as a defensive substance to resist further damage. In turn, Flavonoids together with enzymes in ROS scavenging system scavenge the excess ROS accumulated in vivo to maintain homeostasis.

## DISCUSSION

Dragon’s blood is a precious traditional medicine with multiple therapeutic uses (Masaoud & Schmidt, 1995; Gupta *et al*., 2008; Yang & Yao, 2011; Xin *et al*., 2013; Fan *et al*., 2014; Ding *et al*., 2020; Maděra *et al*., 2020). *D. cochinchinensis* is one of the most important genuine resources of Chinese dragon’s blood as medicine. Here, we constructed its high-quality reference genome at chromosome level combined with PacBio sequencing and Hi-C data. As the first genome report of *Dracaena*, the genome sequence reported herein is expected to contribute to clarify its biology as well as the protection and regeneration of the precious plant resources.

*Dracaena* has been famous all over the world for its longevity (Gupta *et al*., 2008; Zheng *et al*., 2012; Tomlinson & Huggett, 2012; Maděra *et al*., 2018). However, the molecular mechanism underlying its unique biology traits has been unclear. Our analysis revealed *D. cochinchinensis* experienced two genome duplication events, one shared with *A. officinalis* and another independently after their divergence (**Fig. 2b**). Additionally, two LTR insertion burst occurred recently (**Fig. 2c**). Comparative genomic analysis revealed the expanded and unique genes *SAUR* and *cZOGT* are likely the main cause of its longevity and slow growth (**Fig. 2f-g**). Genes in the DNA repair pathway were positively selected which contributes to genome maintenance and environmental adaptability (**Table S13**).

The dragon’s blood, red resin of *Dracaena*, is a defensive substance produced by *Dracaena* when it is damaged or infected by pathogenic microorganisms (Wang, 2007; Jura-Morawiec & Tulik, 2016; Ding *et al*., 2020; Maděra *et al*., 2020). As the main component of dragon’s blood, in addition to pharmacological effects, flavonoids also have many benefits to *Dracaena* itself during the growth development. For example, they are involved in plant defense pathogens and microbes, in absorption of free radicals and ultraviolet light (Neill & Gould, 2003; Crozier *et al*., 2006; Agati *et al*., 2009, 2011; Agati & Tattini, 2010; Landi *et al*., 2015). Our current study focused on analyzing differentially expressed pathways and differential metabolites in the process of dragon’s blood formation, and biosynthesis and regulatory genes in flavonoids biosynthesis in *D. cochinchinensis*. We observed secondary metabolic-related pathways, including flavonoid, terpenoid backbone and phenylpropanoid biosynthesis, were upregulated in 0-6 M after injury (Fig. 4). Correspondingly, metabolome analysis showed the contents of flavonoids compounds are significantly increased, as well as phenylpropanoids, phenols, and phenolic acids. Whereas, the contents of coumarins, organic acids and amines compounds are significantly decreased ((**Fig. 3c**; **Fig. S9; Table S16**). Enrichment in upregulated genes related to biosynthesis of secondary metabolites includes *PAL, CHS*, methyltransferase, etc, which are involved in the flavonoid biosynthesis pathway. Analyzing the dynamic expression profiles of genes in flavonoid biosynthesis pathway during the dragon’s blood formation, showed that most of genes induced expression, even some, such as *CHS, CHI, F3H, FLS* and *PPO*, almost not expressed in health stem, but rapidly and significantly induced after injury, which are considered specifically involved in the formation of dragon’s blood (**Fig. 5**). The expression pattern is essentially consistent with the results we observed differential metabolites and metabolic pathways (**Fig. 3c, Fig. 4**). Further, transcription factors bHLH and MYB are predicted as the core regulators involved in flavonoid biosynthesis pathway (**Figs. S10-12**), which is consistent with previously studies that MYB, bHLH and WD40 TFs are involved in flavonoid metabolism by regulating many structural genes (Hichri *et al*., 2011; Schaart *et al*., 2013). The corresponding relationship between transcription factor and its target was predicted which will provide important clues for studying the function of these genes and their regulation mechanism. Further studies are definitely required to comprehensively elucidate the process of the formation of the red resin in *D. cochinchinensis*, and hence, enable controlled and reasonable dragon’s blood extraction for commercial purposes.

To reveal the regulatory network of dragon’s blood formation induced by injury, we analyzed the signal pathways systematically and considered ROS is the crucial signal stimuli. It is found genes in ROS generation and scavenging are all upregulated after injury (**Fig. S13**), the content of H_2_O_2_ showed regular dynamic changes at different time points after injury (**Fig. S14**), and genes related to the main place of ROS production are expanded (**Table S12, S20**). As we know, *D. cochinchinensis* grows in limestone area where is full of sunlight and strong UV-B, which can cause the degradation of CP43 and CP47 and the decrease of photosynthetic capacity, leads to ROS production and cause DNA damage and protein denaturation (for review Takshak & Agrawal, 2019). KEGG pathway analysis indicated that many of the positively selected genes are related to that, such as base excision repair (BER), nucleotide excision repair (NER), mismatch repair (MMR) and DNA replication (**Table S13**), which implies the *D. cochinchinensis* has a great adaptation to the environment in the evolution to maintain genomic integrity. In addition, genes family in phenylpropanoid biosynthesis and photosynthesis pathway is expanded (**Table S12, S20**). More and more evidences showed flavonoids have multiple photoprotective effects (Winkle-Shirley, 2002; Agati & Tattini, 2010) and functions as scavenger of ROS (Yamasaki *et al*., 1997; Agati *et al*., 2009), as well as moderator of auxin transport (Peer & Murphy, 2007). Taken together, we believe there are some potential associations among “ROS signal--flavonoids biosynthesis--environmental adaptability” in *D. cochinchinensis* (**Fig. 6**). This should be the results of adaptation to the environment in the process of evolution. We speculate that their longevity depends on their strong self-healing ability and adaptability to the environment.

## CONCLUSION

This study reports a chromosome-level genome assembly for *D. cochinchinensis*, the first genome for *Dracaena* Vand. ex L. It provides valuable genetic resources and creates a large scope for studying *Dracaena*. We revealed the genetic basis of its longevity and slow growth at the genomic level, and preliminarily deciphered the molecular mechanism of injury induced flavonoids biosynthesis to form dragon’s blood through transcriptome and metabolome method analysis. These findings are important for an improved understanding of *Dracaena* adaptation, resource protection and utilization and further studies on serving human medicine and health.

## METHODS

### Library Construction and Sequencing

Combined with four types of libraries constructed: (1) the short-read sequencing library with 350-bp insertions from Illumina HiSeq X Ten instrument platforms; (2) The linked read sequencing libraries were constructed on 10X Genomics GemCode platform and sequencing from Illumina platforms; (3) the SMRT Cell libraries with an insert size of 20 Kb for PacBio sequencing; and (4) the chromatin interaction mapping (Hi-C) library with 350-bp insertions for Illumina sequencing.

### Genome Assembly and Assessment of the Assembly Quality

*De novo* assembly of the PacBio single-molecule long reads from SMRT Sequencing was performed using FALCON (https://github.com/PacificBiosciences/FALCON/) (Chin *et al*., 2013). BWA-MEM (Li *et al*., 2013) was used to align the 10X Genomics data to the assembly using default settings. Scaffolding was performed by fragScaff (version 1.1) with the barcoded sequencing reads. These contigs were then used to form super-scaffolds. BWA (v0.7.8) (Li & Durbin, 2009) software was used to align the Hi-C clean data to the preceding assembly. According to the linkage information and restriction enzyme site, the string graph formulation was used to construct the scaffold graph with LACHESIS (Burton *et al*., 2013).

### Genome Annotation

Genome repeat sequences were annotated de novo by using RepeatMasker (http://www.repeatmasker.org). The protein-coding genes were predicted based on the repeat-masked genome, conserved protein homologues and assembled transcripts. miRNA and snRNA genes were predicted by INFERNAL software using the Rfam database (Supplementary method).

### Repetitive Sequences Analysis

TEs identification was performed using a combination of homology-based and ab initio-based approaches. For the homology-based method, RepeatMasker (v3.3.0) was applied to identify the TEs with a known repeat library (Repbase 15.02). For the ab initio-based approach, TEs prediction was implemented by RepeatScout (Price *et al*., 2005), RepeatModeler (v1.0.5; http://www.repeatmasker.org/RepeatModeler/), Piler, and LTR_FINDER (Xu *et al*., 2007) to build the de novo repeat libraries. RepeatMasker (v3.3.0) (Chen, 2004.) was run on the *D. cochinchinensis* genome assembly using the de novo libraries as reference libraries. Tandem Repeats Finder (TRF) (Benson, 1999) was used to identify the tandem repeats with integrated repeat library.

We further investigated the dynamics of long terminal repeats retrotransposons (LTR-RTs) in the *D. cochinchinensis* genome. *De novo* detection of LTR-RTs was performed using LTRharvest (-motif tgca -motifmis 1) and LTRfinder (-minlenltr 100 -maxlenltr 5000 -mindistltr 1000 -maxdistltr 20000). The primer binding site (PBS) motifs were annotated using LTRdigest, and only elements containing PBS motifs were retained for further domain annotation. Domains of retrotransposons were identified by searching against HMM profiles collected by Gypsy Database (http://gydb.org/), and internal features of LTR-RTs were annotated by LTRdigest. Intact LTR-RTs that contained domains (gag domain, protease domain, reverse transcriptase domain and integrase domain) were clustered, and those belonged to families with less than 5 members were discarded. Nucleotide divergence rate (λ) was calculated from the MUSLCE alignment of the 5′and 3′LTR sequences of LTR-RTs by EMBOSS program distmat. The insertion date (T) was computed for each LTR-RT (T = K/2r, K: genetic distance, K = − 0.75ln (1 − 4λ/3)), which was set substitution rate (r) of 1.3 × 10− 8 substitutions per site per year.

### Gene Family Construction and Phylogenetic Analysis

Gene families were generated by OrthoMCL (http://orthomcl.org/orthomcl/) (Li *et al*., 2003) from 17 species, including *Dracaena cochinchinensis, Asaragus officinalis, Dendrobium officinale, Apostasia shenzhenica, Elaeis guineensis, Oryza sativa* (phytozomev10), *Ananas comosus, Musa acuminate, Arabidopsis thaliana* (phytozomev10), *Populus trichocarpa, Aqularia sinensis, Hevea brasiliensis, Citrus grandis, Carica papaya, Eucalyptus grandis, Vitis vinifera* (phytozomev10), *Amborella trichopoda* with the parameter of 1.5 inflation index.

Protein sequences from 252 shared single-copy gene were utilized to construct the phylogenetic tree. Protein sequences of these orthologs were aligned by MUSCLE (Edgar, 2004), and the phylogenetic tree was constructed by the ML (maximum likelihood) TREE algorithm in RAxML software (Stamatakis, 2006). Then mcmctree program of PAML (http://abacus.gene.ucl.ac.uk/software/paml.html) (Yang, 2007) was applied to estimate divergence time among 17 species. Five calibration points were selected and applied in the present study: *V. vinifera and O. sativa* time (130∼200 Mya) (Iorizzo, et al., 2016), *O. sativa and A. shenzhenica* time (137∼156 Mya) (Zhang *et al*., 2017), *V. vinifera and C. papaya* time (114∼134 Mya) (Ming *et al*., 2008), *P. trichocarpa and A. thaliana* time (107∼109 Mya) (Tuskan *et al*., 2006), *H. brasiliensis* and *P. trichocarpa* time (65∼86 Mya) (Tang *et al*., 2016; Tuskan *et al*., 2006). Gene gain and loss in gene families were determined using CAFÉ (v2.2) (Computational Analysis of gene Family Evolution). The program uses a birth and death process to model gene gain and loss over a phylogeny.

### Expansion and Contraction of Gene Families

Gene gain and loss in gene families were determined using CAFÉ (v2.2) (Computational Analysis of gene Family Evolution). The program uses a birth and death process to model gene gain and loss over a phylogeny. Enrichment of KEGG pathway for expanded gene families of *D. cochinchinensis* was summarized and visualized using KOBAS software.

### RNA Sequencing and Analysis

Five tissues (root, stem, leaves, flower, fruit) were harvested from *D. cochinchinensis*, and 33 cut injury-treated stems at different time points (0h, 2h, 6h, 12h, 24h, 2d, 3d, 4d, 5d, 10d, 15d, 30d, 6M, 17M). Three biological replicates for each tissue were collected. Raw reads from different samples were filtered to obtain high-quality clean reads following standard process. For gene expression analysis, clean reads were mapped to *D. cochinchinensis* genome using Tophat (v2.0.8) (Kim *et al*., 2013). Then the readcounts of each gene was normalized for downstream analysis. A gene is defined as expressed if its RPKM value is ≥1 in at least one of all the transcriptomes.

Differential expression analysis between a treatment sample and the control sample was performed using the DEseq R package (v1.18.0). The resulting P-values were adjusted using the Benjamini and Hochberg’s approach for controlling the false discovery rate. Genes with an adjusted P-value <0.05 found by DESeq were assigned as differentially expressed. KEGG pathway enrichment analysis of the DEGs was performed using KOBAS software (Mao *et al*., 2005).

### Metabolic Profiling Acquisition Based On UHPLC-Q-Orbitrap-MS

The freeze-dried samples were dissolved in 50% methanol-water (*v/v)* solutions at the concentration of 5 g·mL^-1^, and the solutions were filtered, centrifuged at 13,200 × *g* for 10 min, and 10 μL of the resulting supernatant was directly injected into the LC-MS system.

Untargeted metabolomics was accomplished using a U3000 UHPLC equipped with aQ-Exactive™ Q-Orbitrap MS through a HESI source (Thermo Fisher Scientific, Waltham, MA, USA). A Waters Acquity UPLC HSS T_3_ column (2.1 × 100 mm, 1.8 μm) was used for chromatographic separation with a column temperature of 35°C. The flow rate was set as 0.4 mL·min^-1^. The autosampler temperature was maintained at 4°C.

The mobile A and B phases were 0.1% formic acid–water (*v/v*) and acetonitrile, respectively. The following elution gradient was used: 10% (B) in 0–3 min, 10–22% (B) in 3-5 min, 22-27% (B) in 5-7 min, 27-45% (B) in 7-25 min, 45-57% (B) in 25-35 min, 57-100% (B) in 35-36 min, 100% (B) in 36-37 min, 100-10% (B) in 37-37.5 min, and 10% (B) in 37.5-40 min.

The HESI source parameters were as follows: the scan mode was full-ms/dd-ms mode and the scanning range was *m/z* 150-1,500 at a resolution of 70,000; the spray voltage was 3.0 kV in the positive mode and 2.8 kV in negative mode; the capillary temperature and the probe heater temperatures were 320°C and 350°C, respectively; the sheath gas (N_2_) was 35 arbitrary units (arb) and the auxiliary gas was 10 arb (99.9% pure N_2_). The S-lens level value was 50 V. The normalized collision energy was 20/40/60 V. Data acquisition was conducted using Xcalibur 4.1 software (Thermo Fisher Scientific).

## FOUNDS

This work was supported by the program of CAMS Initiative for Innovative Medicine (No. 2021-1-I2M-032); the National Natural Science Foundation of China (82173925, 81573525); the National Key Research and Development Program of China (Grant NO. 2018YFC1706401).

## Author Contributions

YX and JW conceived and designed the project, and participated in all the related analyses. YX and KZ performed the data analyses and drafted the manuscript. KZ performed genome assembly and annotated the genome. ZZ conducted metabolome analysis. YL conducted H_2_O_2_ analysis. YX, ZZ and YL contributed to the sample preparation used for genome and RNA sequencing. All authors contributed to the writing of the paper.

## Data Availability Statement

The raw data of genome and transcriptome sequencing reported in this paper will be deposited in the Genome Sequence Archive (Wang et al., 2017) in BIG Data Center, Beijing Institute of Genomics (BIG), Chinese Academy of Sciences under the accession number (PRJCA007701). It will be publicly accessible at http://bigd.big.ac.cn/gsa) when paper is accepted.

## ACKNOWLEDGMENTS

The authors declare no competing financial interests

## Notes

### Competing Interest Statement

The authors have declared no competing interest.

