## supplemental methods for "The Chromosome-level Genome of *Dracaena cochinchinesis* Provides Insights into its Biological Features and the Mechanism of Dragon’s Blood Formation"

**Methods S1. Genome size estimation**

44.11 G sequencing data of *D. cochinchinensis* genome quality-filtered reads from the Illumina platform were subjected to 17-mer frequency distribution analysis with SOAPdenovo (Luo *et al*., 2012). K-mer 17 was selected to estimate the genome size and heterozygosity of *D. cochinchinensis*. We plotted the distribution of k-mer depth against frequency with a main peak occurred at the depth of 28. Based on the total number of k-mers, The *D. cochinchinensis* genome size was calculated using the following formula: genome size = k-mer_Number/ Peak_Depth and Revised Gsize=Genome size × (1-Error Rate).

**Methods S2. Library construction and sequencing**

Total genomic DNA was isolated from young leaves of the *D. cochinchinensis* using Plant DNA Kit (TIANGEN) according to the manufacturer’s instructions. The DNA was sheared by Covaris® M220 focused-ultrasonicator^TM^ instrument (Covaris, Woburn, Massachusetts, USA). The sheared DNA, with fragment sizes of 350 bp, was processed using the TrueSeq DNA PCR-Free LT Library Kit protocol. This PCR-Free library was sequenced with a HiSeq X Ten instrument as 150 bp PE reads.

40 μg of sheared DNA was used to construct SMRT Cell libraries with an insert size of 20 Kb. The libraries were then sequenced with a PacBio sequel instrument (Pacific Biosciences, Menlo 31 Park, CA, USA). The linked read sequencing libraries were constructed on 10X Genomics GemCode platform. Sample indexing and partition barcoded libraries were prepared using the Chromium Genome Reagent Kit (10x Genomics) according to manufacturer’s instruction. The barcode sequencing library was firstly quantified by Qubit2.0, insert-size was checked by Agilent2100, and finally quantified by qPCR. The 130.40 Gb library sequenced with 150bp paire-end reads on an Illumina HiSeq X Ten platform.

For the Hi-C library, chromatin was fixed in place with formaldehyde in the nucleus. Fixed chromatin was digested with DpnII restriction endonuclease, 5′ overhangs filled in with biotinylated nucleotides, and free blunt ends were ligated. After ligation, cross-links were reversed and the DNA was purified from protein. Purified DNA was treated to remove biotin that was not internal to the ligated fragments. The DNA was then sheared to a mean fragment size of ~350 bp, and sequencing libraries were generated using NEBNext Ultra enzymes and Illumina-compatible adaptors. Biotin-containing fragments were isolated using streptavidin beads before PCR enrichment of each library. The libraries were sequenced on an Illumina HiSeq platform.

**Methods S3. Genome assembly and assessment of the assembly quality**

*De novo* assembly of the PacBio single-molecule long reads from SMRT Sequencing was performed using FALCON (<https://github.com/PacificBiosciences/>FALCON/). Before assembly, we used FALCON to correct the PacBio reads and then assemble them into contigs with parameters (length_cutoff_pr = 5000, max_diff = 120, max_cov = 130). The draft assembly was polished using the quiver algorithm. Then, Pilon (Walker *et al*., 2014) were used to perform error correction of p-contigs with 102.92X coverage of short paired-end reads generated from Illumina HiSeq Platforms.

We used BWA-MEM (Li *et al*., 2013) to align the 10X Genomics data to the assembly using default settings. Scaffolding was performed by fragScaff (version 1.1) with the barcoded sequencing reads. These contigs were then used to form super-scaffolds. BWA (v0.7.8) (Li & Durbin, 2009) software was used to align the Hi-C clean data to the preceding assembly. Only the read pairs with both reads aligned to contigs are considered for scaffolding. The mis-assembly can be detected by the sudden drop in per-base physical coverage in a contig. According to physical coverage of the alignment result, the mis-assemblies will be sheared to correct the misassemble errors by SALSA (Ghurye *et al*., 2017). According to the linkage information and restriction enzyme site, the string graph formulation was used to construct the scaffold graph with LACHESIS (Burton *et al*., 2013).

The *D. cochinchinensis* assembly was further refined using ~137.26 Gb Hi-C data, 1,094 Mb (90.26%) of the contig sequences were anchored onto 20 chromosomes. The scaffold N50 was finally improved to be 50.06 Mb, with the longest scaffold being 114.845 Mb.

**Methods S4. Gene annotation**

Genes of the *D. cochinchinensis* genome were annotated using multiple methods, including homology-based predictions, *de novo* predictions and transcriptome-based predictions. For *de novo* predictions, Augustus (v3.0.2) (Stanke *et al*., 2006), GENSCAN (v1.0) (Burge & Karlin, 1997), GlimmerHMM (v3.0.2) (Majoros *et al*., 2004), Geneid (v1.4) (Parra *et al*., 2000) and SNAP (v11-29-2013) (Korf, 2004) analysis were performed on the repeat-masked genome. Predicted protein sequences from *Oryza sativa* (phytozomev10), *Elaeis* *guineensis* (Singh *et al*., 2013), *Asaragus* *officinalis* (Harkess *et al*., 2017), *Ananas* *comosus* (Ming *et al*., 2015), *Dendrobium* *officinale* (Yan *et al*., 2015), *Musa* *acuminata* (D’hont *et al*., 2012), *Arabidopsis* *thaliana* (phytozomev10) were used for homology-based predictions. First, query sequences were subjected to TBLASTN analysis with an Expect (E)-value cutoff of 1e-5. The homologous genome sequences were aligned against the matching proteins using GeneWise software (v2.2.0) (Birney *et al*., 2004) for accurate spliced alignments. For transcriptome-based predictions, RNA-seq data were used for gene annotation processed by TopHat (v2.0.8) and Cufflinks (v2.1.1) (Trapnell *et al*., 2009).). RNA-seq data were also assembled by Trinity (Haas *et al*., 2013). PASA software (v2.3.3) (Haas *et al*., 2003) was then performed to generate a full transcriptome-based genome annotation. The homology, *de novo* and transcriptomic gene sets were merged to form a comprehensive and non-redundant reference gene set using EVidenceModeler (v1.1.1) (Haas *et al*., 2008) software. Next, PASA (v2.3.3) (Haas *et al*., 2003) was used to generate UTRs, alternative splicing variation information.

Functional annotation of the protein-coding genes was carried out using blastp (E-value cut-off 1e**-**05) against SwissProt (http://www.uniprot.org/) (Bairoch & Apweiler, 2000) and NR database. Protein domains were annotated by searching against InterPro (v29.0) (Mulder & Apweiler, 2007) and Pfam database (Finn *et al*., 2014), using InterProScan (v4.7) (Mulder & Apweiler, 2007) and HMMER (http://hmmer.janelia.org), respectively. The GO terms for genes were obtained from the corresponding InterPro or Pfam entry. The pathways in which the genes might be involved were assigned by BLAST against the KEGG database (release 53) (Kanehisa & Goto, 2000) with the E-value cut-off of 1e-05.

**Methods S5. Non-coding RNA annotation**

The annotation of tRNA was performed using tRNAscan-SE (Lowe & Eddy, 1997) software with default parameters. rRNA annotation was based on homology with rRNAs from several diverse higher plant species (not shown), using blastn with ‘E-value = 1e-5’. miRNA and snRNA genes were predicted by INFERNAL software (Nawrocki et al., 2009) using the Rfam database (Griffiths-Jones et al., 2005).

**Methods S6. Positively selected genes in *D. cochinchinensis***

The protein alignments of single-copy gene families in positive species were generated using MUSCLE (Edgar, 2004). Gblocks (Castresana, 2000) was applied to filter poorly aligned positions and divergent regions of the protein alignments before transformed to CDS alignments. With the foreground branch, positive selection sites were detected based on branch-site models of PAML (v4.9) (Yang, 2007) using the CDS alignments. P-values were computed using the χ^2^ statistic and adjusted by FDR method. Finally, positively selected genes were found in *Dracaena cochinchinensis* genome.

**Methods S7. Microscopic Observation**

Microscopic observation was made after 1, 2, and 3 months of wounding stress to view the formation of dragon's blood. The detailed methods were as follows: approximately 2 cm3 (depth: approximately 1 cm) wood slices were collected from the wounding locations, and the wood slices were cut into blocks (3–5 mm2) as thin as possible with a razor blade. The blocks were treated with chloral hydrate penetration solution and dilute glycerol, and then observed and photographed using a microscope (DXM1200C; Nikon, Tokyo, Japan) and camera (ECLIPSE 80i, Nikon, Tokyo, Japan).

**Methods S8. Data processing of metabolomic assay**

Data processing of the metabolomic assays was performed using SIEVE 2.2 (Thermo Fisher Scientific) for background subtraction and component extractions. The obtained peak list was further processed by principal component analysis (PCA) and Student's *t*-tests. PCA was performed by the SIMCA-P program (version 14.1 (Umetrics, Umea, Sweden). Student's *t*-test was conducted using Office Excel 2010 (Microsoft, Redmond, WA, USA). SIMCA-P 14.1 was also used for data transformation for orthogonal partial least squares discriminant analysis (OPLS-DA). Metabolites satisfying both VIP > 1.0 and *p* < 0.01 were chosen concurrently for further screening for differential compounds before and after wounding stress. The metabolites were identified or tentatively identified using the following: (1) Metlin database (http://metlin.scripps.edu/index.php), *m/z* cloud (https://www.mzcloud.org/), HMDB (http://www.hmdb.ca/), and Compound Discoverer 2.1 (Thermo Fisher Scientific); (2) A self-constructed LC-MS/MS identification system for metabolites; and (3) Comparison with reference compounds derived from *Dracaena* plants and dragon's blood. Additionally, Multiple Experiment Viewer (MEV) software (<http://mev.tm4.org/>) was used for the generation of heat maps. Prism 6 (GraphPad, San Diego, CA, USA) was used for producing box plots and line graphs.

**Methods S9. H_2_O_2_ change after mechanical damage**

0.5 cm-diameter holes are drilled in the stem of six-year-old *D. cochinchinensis* trees to induce the mechanical dagame. 0.8x0.8 cm^2^ stem blocks containing the hole are taken 0.5 h,1 h, 2 h, 3 h, 4 h , 6 h, 8 h,12 h, 16 h, 20 h, 24 h, 2 d, 3 d, 5 d, and 10 d after damage. Stem blocks without hole is taken as negative control. Stem blocks are powdered with liquid nitrogen. 0.1 g powder is fully mixed into 400uL 4°C pre-cooling acetone and then is centrifuged by 3000 rmp for 20 min under 4°C. Collect the supernatant. Then 200 uL 4°C pre-cooling acetone is added into the sediment. The mix is centrifuged by 3000rmp for 20min under 4°C. Merge the supernatant as the H_2_O_2_ extract. The H_2_O_2_ concentration is detected by the ELISA kit following the introduction. Statistical analysis is calculated using SPSS software.
