## supplemental figures for "The Chromosome-level Genome of *Dracaena cochinchinesis* Provides Insights into its Biological Features and the Mechanism of Dragon’s Blood Formation"

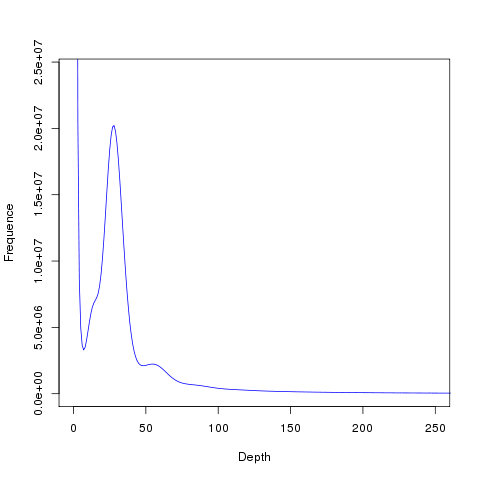

**Supplemental Figure 1.** Frequency Distribution of K-mer Depth and K-mer Numbers (K-mer = 17)

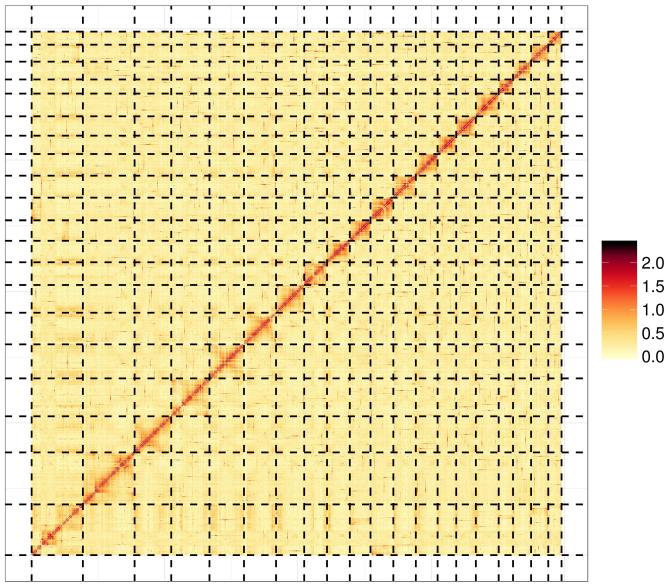

**Supplemental Figure 2.** Chromosome-level assembly of the *D. cochinchinensis* genome using Hi-C technology.

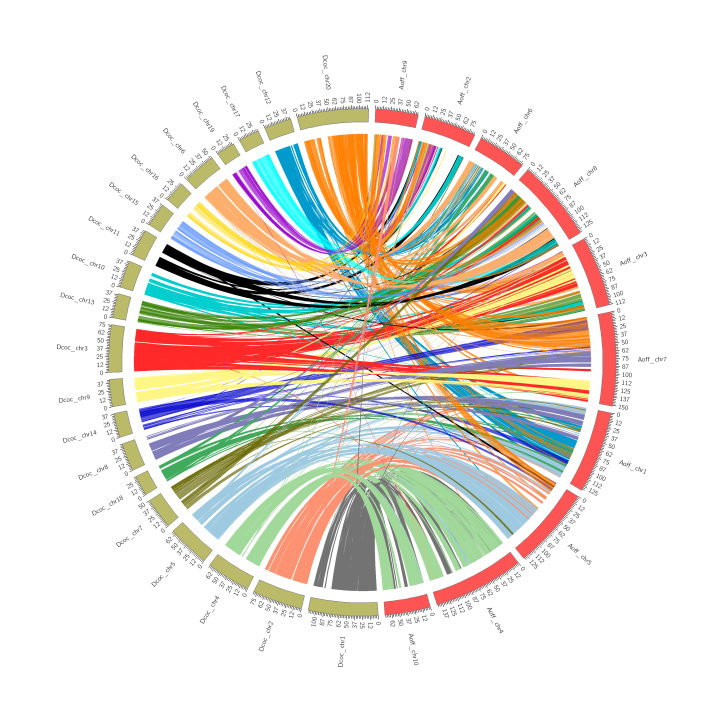

**Supplemental Figure 3.** Synteny between *D. cochinchinensis* and *A. officinalis* Chromosome.

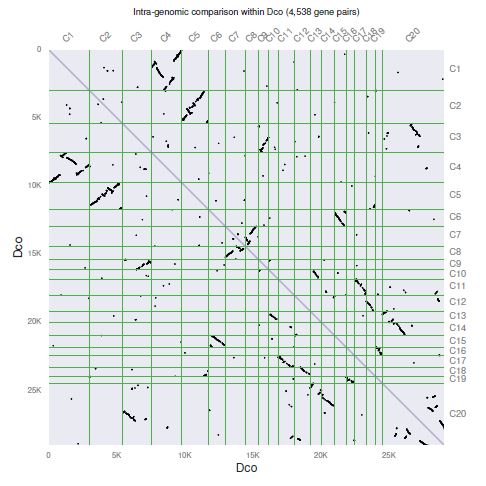

**Supplemental Figure 4.** Intra-genomic comparison within *D. cochinchinensis*.

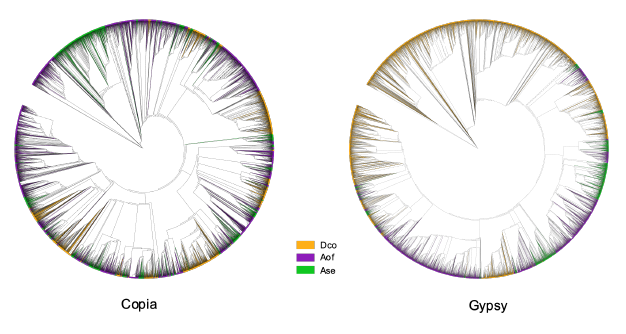

**Supplemental Figure 5.** Phylogenetic analysis of reverse transcriptase (RT) genes from complete retrotransposons in *D. cochinchinensis*, *A. officinalis* and *A. shenzhenica*.

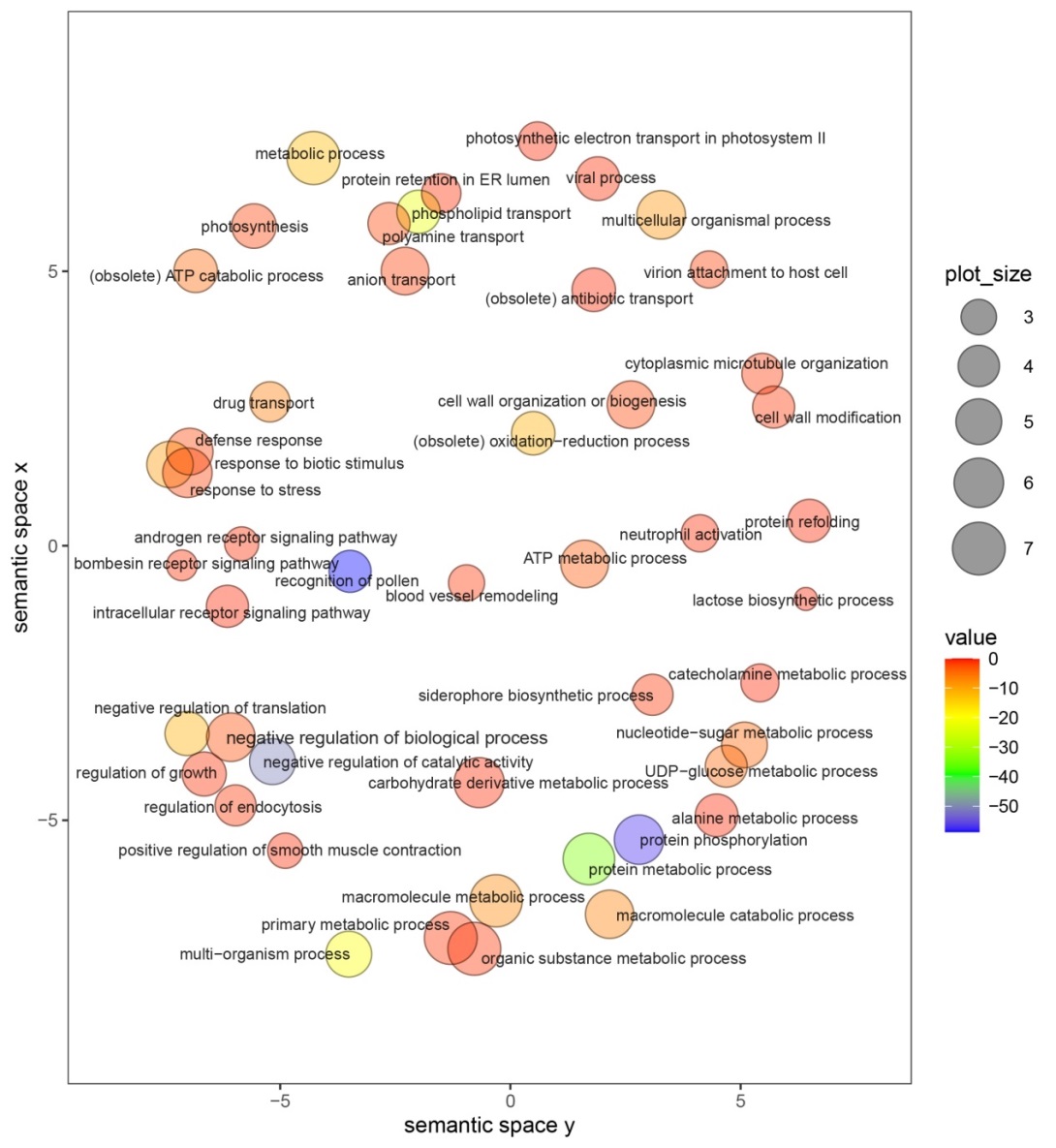

**Supplemental Figure 6.** **REViGO semantic similarity scatter plot of Biology Process Gene Ontology terms for expanded genes in *D*. *cochinchinesis*.** In semantic spaces, the proximity between circles represents relatedness (similarity) of the GO terms. Similar GO terms are close together in the plot. The axes in the plot have no intrinsic meaning, but were used to measure pairwise similarities between GO terms. Color indicates degree of enrichment for each process presented as the p-value from the hyper-geometric test.

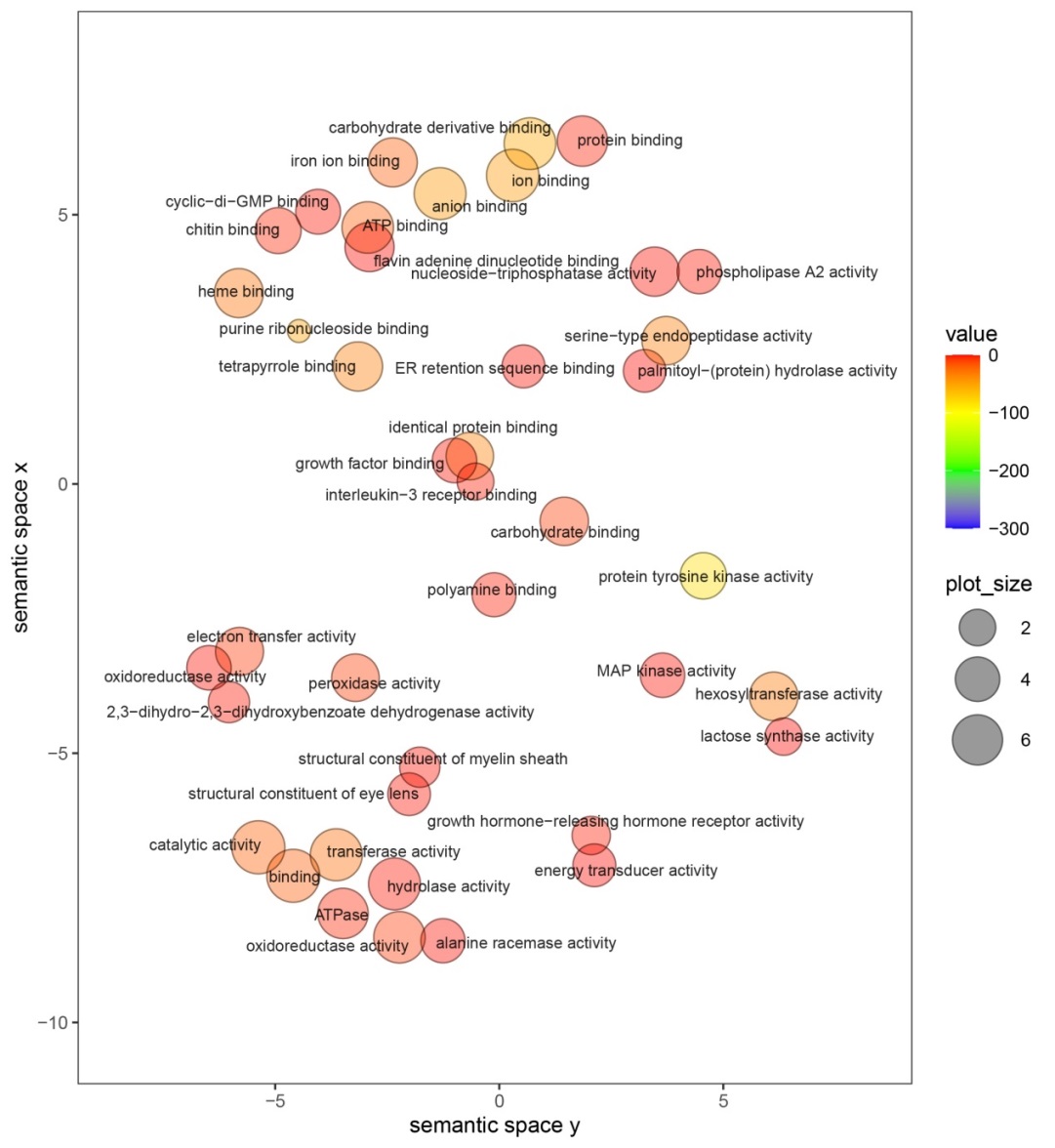

**Supplemental Figure 7. REViGO semantic similarity scatter plot of Molecular Function Gene Ontology terms for expanded genes in *D*. *cochinchinesis*.** In semantic spaces, the proximity between circles represents relatedness (similarity) of the GO terms. Similar GO terms are close together in the plot. The axes in the plot have no intrinsic meaning, but were used to measure pairwise similarities between GO terms. Color indicates degree of enrichment for each process presented as the p-value from the hyper-geometric test.

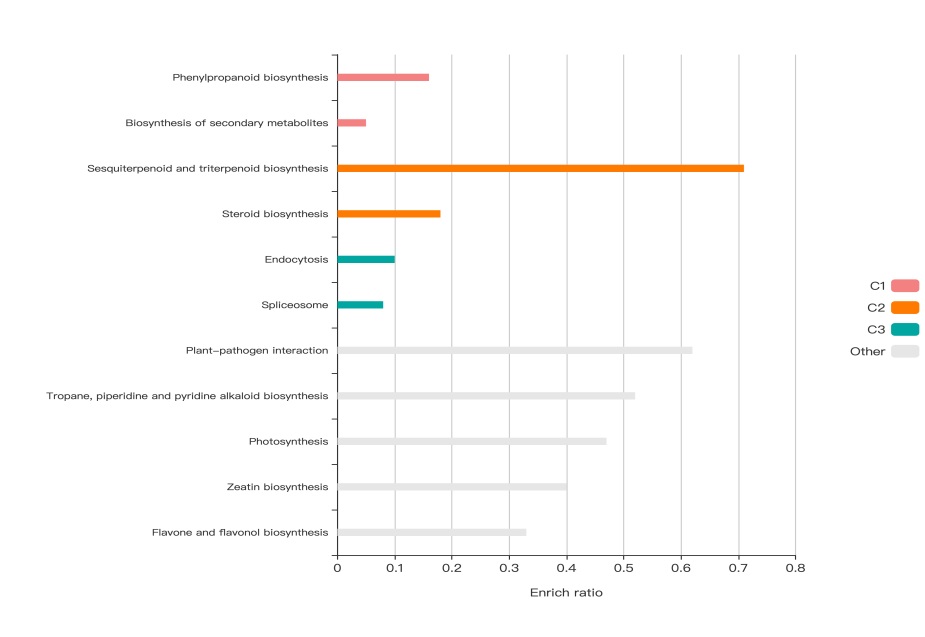

**Supplemental Figure 8. KEGG enrichment barplot of expanded genes in *D*. *cochinchinesis*.**

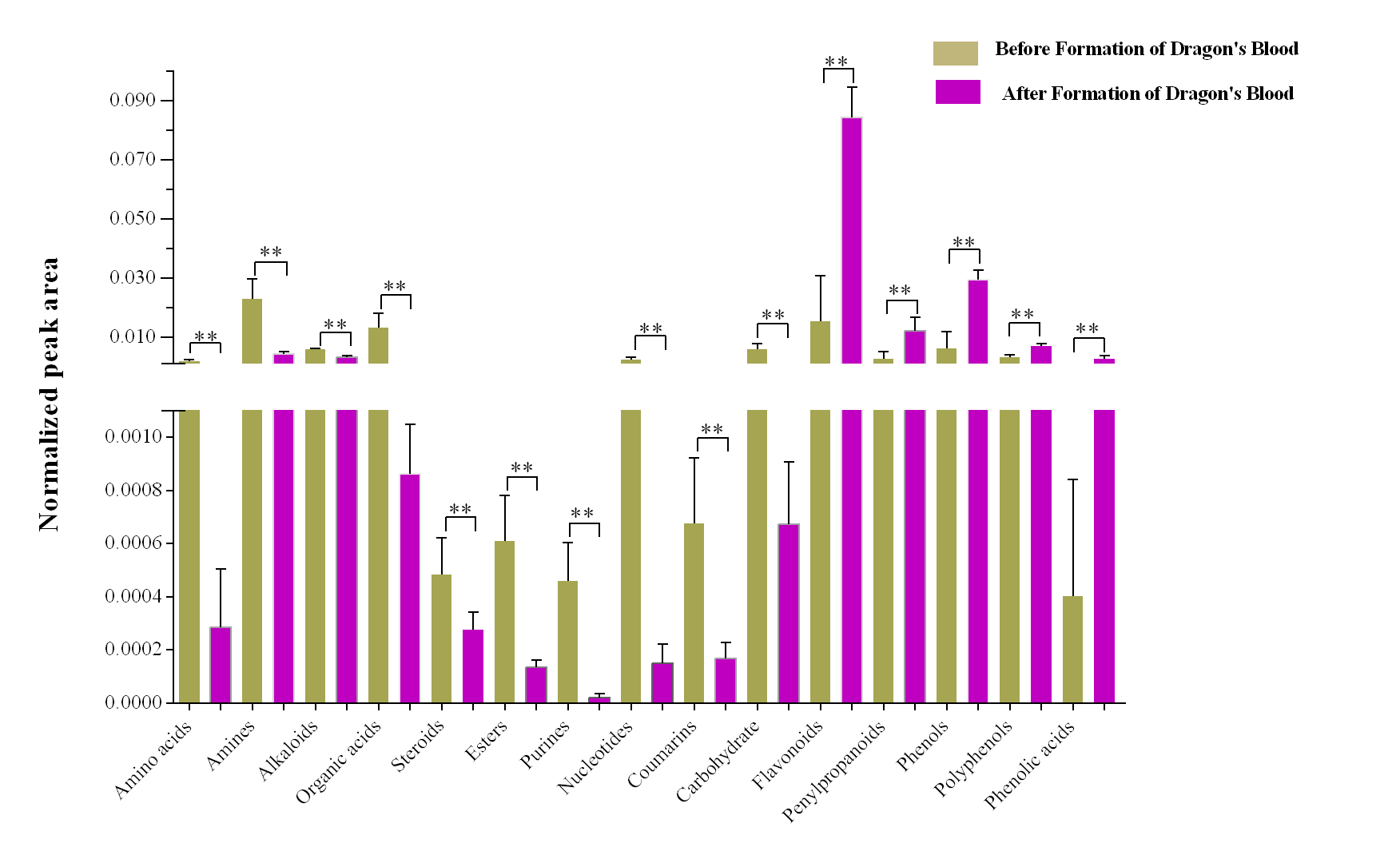

**Supplemental Figure 9. Histogram of the normalized peak area of metabolite categories before and after the wounding treatment.** (**: p < 0.01.)

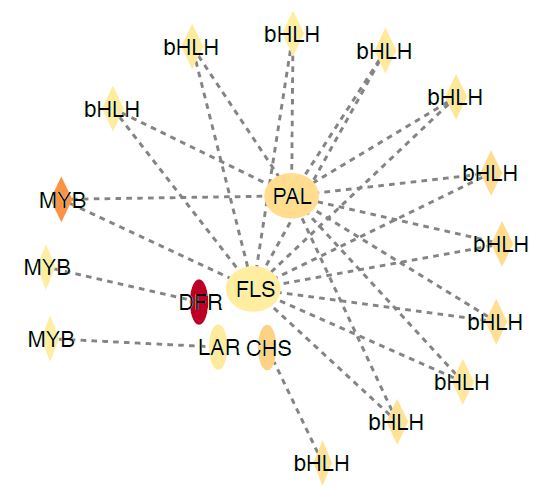

**Supplemental Figure 10. Co-expression analyses of DEG genes showed transcription factors bHLH and MYB are the core regulators of flavonoids biosynthesis*.***

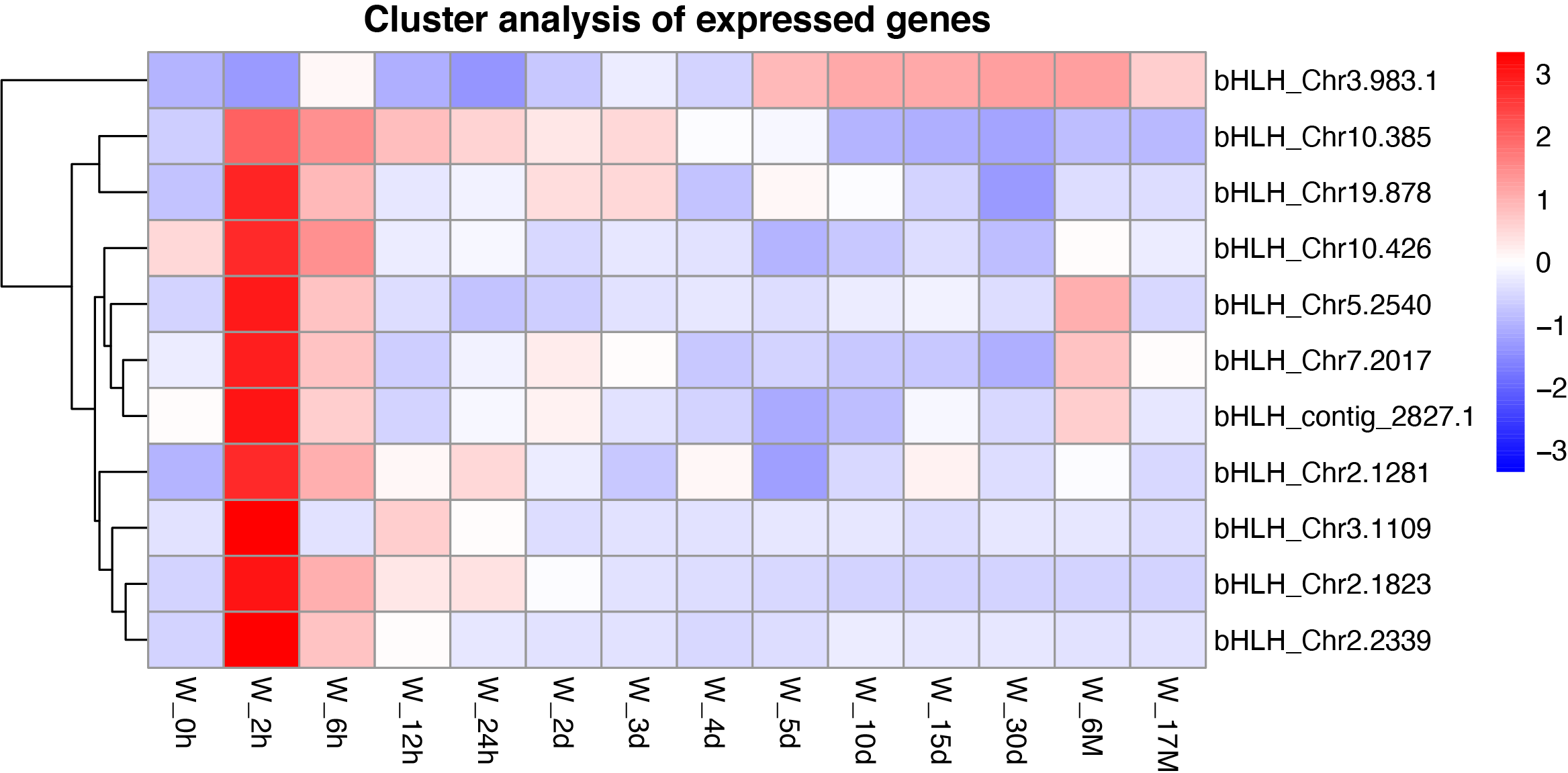

**Supplemental Figure 11. Heatmap of transcription factors bHLH predicted to regulate the flavonoids biosynthesis pathway.**

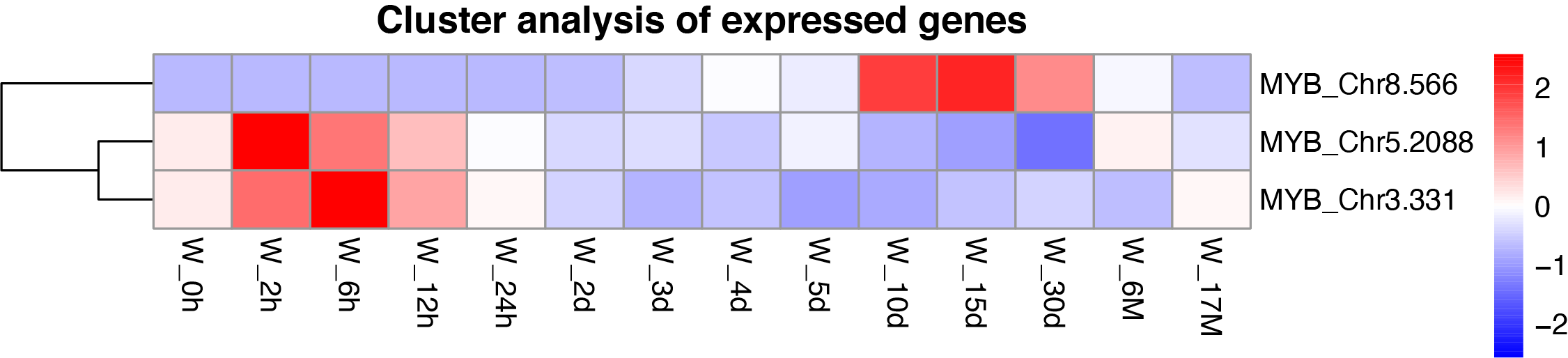

**Supplemental Figure 12. Heatmap of transcription factors MYB predicted to regulate the flavonoids biosynthesis pathway.**

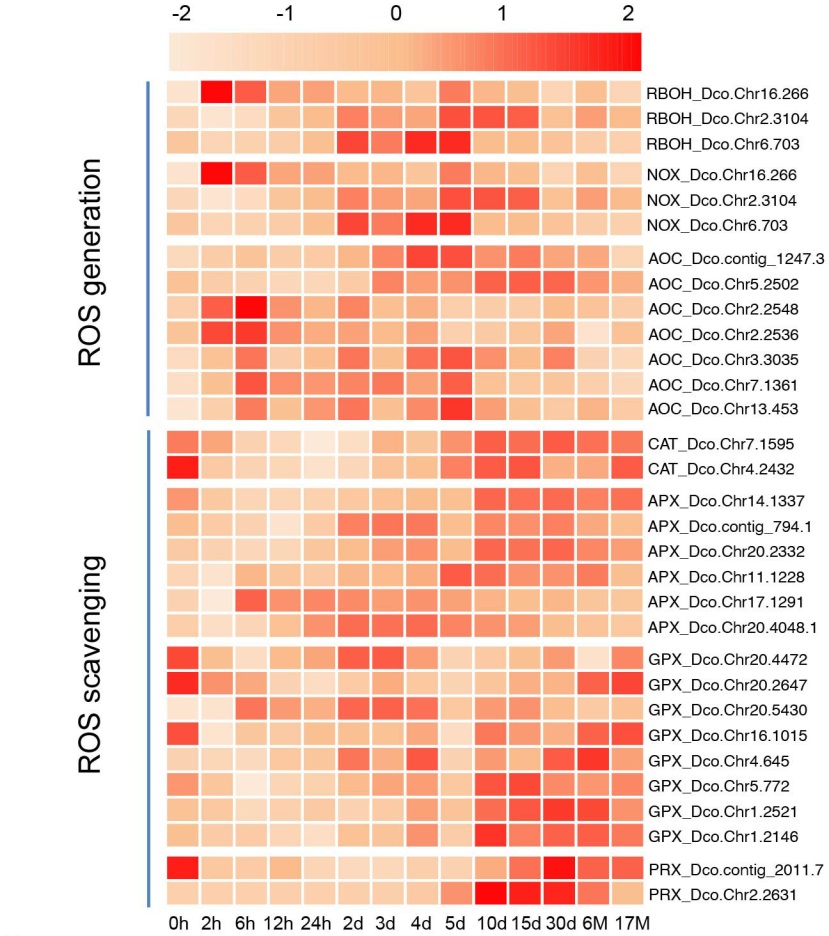

**Supplemental Figure 13. Heat map of genes involved in ROS generation and scavenging.** RBOH: respiratory burst oxidase homolog; NOX: NAD(P)H oxidases; AOC: amine oxidases. CAT: catalase; APX: ascorbate peroxidase; GPX: glutathione peroxidase; PRX: peroxidase.

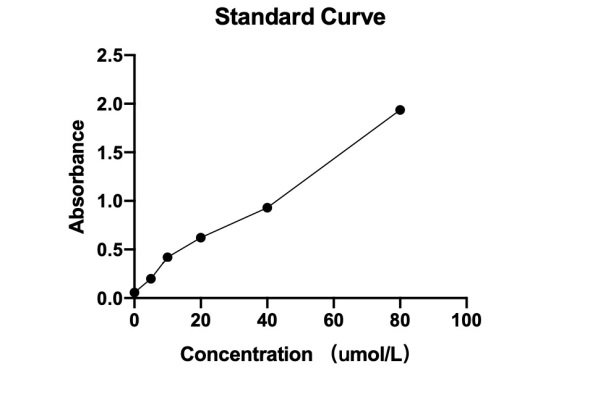

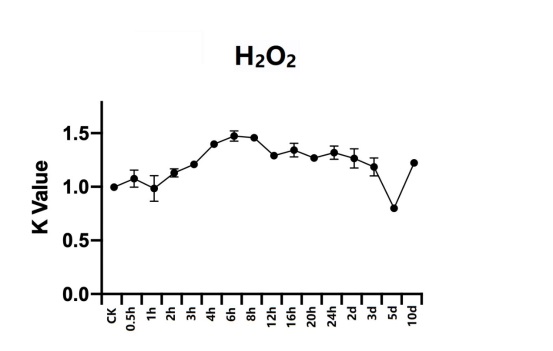

**Supplemental Figure 14. H_2_O_2_ Concentration Change over time after Mechanical Damage.** a. standard curve; b. dynamic change of H_2_O_2_ concentration.

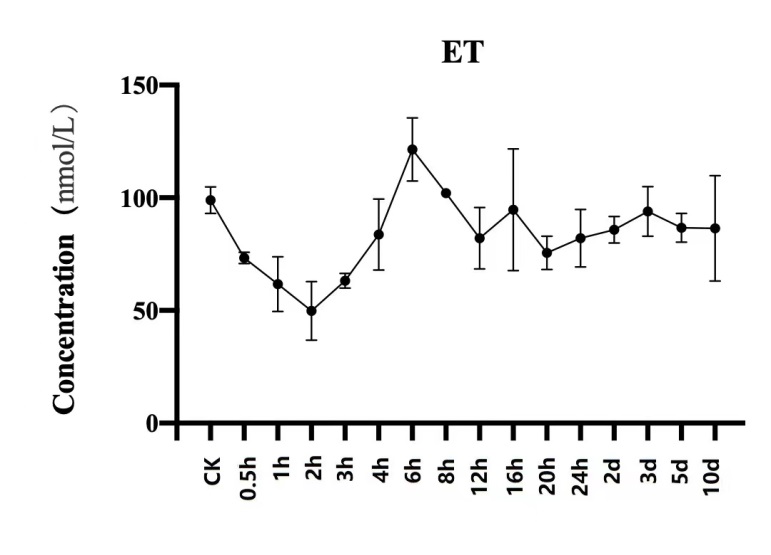

**Supplemental Figure 15.** **Ethylene (ET) Concentration Change over time after Mechanical Damage.**

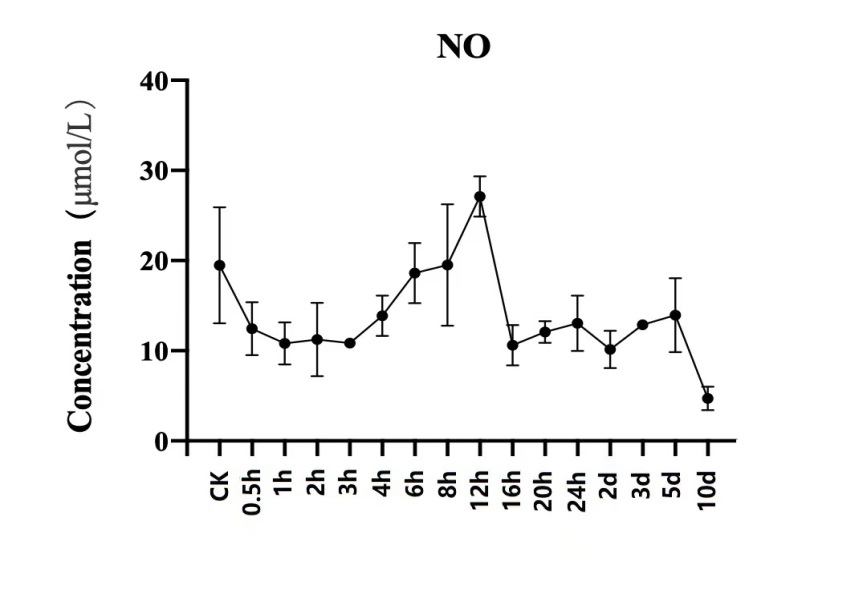

**Supplemental Figure 16.** **Nitric Oxide (NO) Concentration Change over time after Mechanical Damage.**

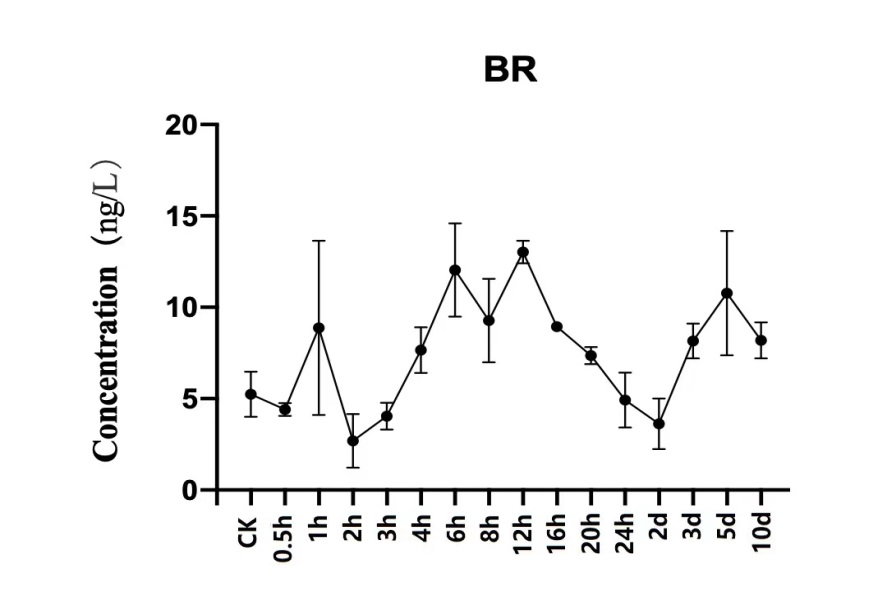

**Supplemental Figure 17.** **Brassinosteroids (BR) Concentration Change over time after Mechanical Damage.**
